# Release of the *Staphylococcus aureus* autoinducing peptide is enabled by active export of the biosynthetic intermediate

**DOI:** 10.64898/2026.09.29.755489

**Authors:** Alan Tong, Kelsey M. Hallinen, Arya Bhakare, Steven P. Bodine, Zemer Gitai, Tom W. Muir

**Affiliations:** Department of Chemistry, Princeton University, Princeton, NJ 08544; Department of Physics, Princeton University, Princeton, NJ 08544; Department of Physics, University of Oregon, Eugene, OR 97403; Department of Chemistry, Stanford University, Stanford, CA 94305; Department of Molecular Biology, Princeton University, Princeton, NJ 08544

**Keywords:** Quorum sensing, Transporters, *Staphylococcus aureus*, Autoinducing peptide

## Abstract

*Staphylococcus aureus* is a species of Gram-positive bacteria that uses the accessory gene regulator (*agr*) quorum sensing system to regulate many aspects of cell behavior such as virulence and toxin production. Quorum sensing is activated in dense *S. aureus* populations by the extracellular accumulation of a signaling molecule called the autoinducing peptide (AIP). While the biosynthetic enzymes that produce AIP have been elucidated in certain subgroups of *S. aureus*, the mechanism of AIP secretion is unknown. Here, we show that the biosynthetic intermediate known as AgrD (1-32) thiolactone is actively exported into the surrounding environment in certain *S. aureus* subgroups. Using a high-throughput transposon mutagenesis screen, we identify the integral membrane transporters, AbcA and PmtCD, as being involved in the biosynthesis of certain AIPs. Subsequent genetic complementation and biochemical experiments reveal that these proteins actively transport AgrD (1-32) thiolactone out of the cell. This enables the production of AIP directly on the cell surface following the action of MroQ, an integral membrane protease demonstrated here to possess an active site facing the extracellular milieu. Furthermore, we show that media supplementation with catalytically-active MroQ nanodiscs rescues AIP production in an *mroQ* knockout strain. This observation suggests the possibility that some proportion of AgrD (1-32) thiolactone exported by one cell may travel to neighboring cell surfaces for further processing into the diffusive AIP molecule. Collectively, our studies shed light on a poorly understood step in the *agr* quorum sensing pathway with clear implications for the design of novel therapeutic strategies.

## Introduction

Quorum sensing (QS) is the process in which a single bacterium can specifically detect and react to the concentration of autoinducer molecules, which act as local indicators of cell density (1). Once the bacterium senses that the autoinducer concentration is above a certain threshold, there is a change in the regulation of various gene expression programs that enables the cell to contribute to synchronized group behaviors. Among Gram-positive bacteria, the QS system of *Staphylococcus aureus* serves as an important model system. *S. aureus* is an opportunistic human pathogen known for its potential to develop resistance against all classes of clinically available antibiotics, and methicillin-resistance *S. aureus* (MRSA) is a particularly notable example of a superbug strain (2). The *S. aureus* QS system is encoded in a genomic site called the accessory gene regulator (*agr*) locus, which encodes four genes (*agrBDCA*) under the control of the P2 promoter (3, 4). This system is regulated by a cyclic pheromone known as the autoinducing peptide (AIP) (5). Low AIP concentrations are associated with biofilm formation and increased expression of defensive proteins such as protein A (a cell-surface protein involved in immune evasion) and coagulase (a protective fibrin coat-forming enzyme) (6–8). Once the bacterial cells grow past a threshold density, the high concentration of AIP triggers biofilm dispersal and increased expression of virulence factors such as hemolysins and enterotoxins (9, 10). Thus, the accumulation of AIP causes a complex yet coordinated defense-to-offense lifestyle change in *S. aureus*, a transition that is critical for its extreme pathogenicity during infections. Elucidating how AIP is biosynthesized, the mechanism by which it is secreted, and the genetic factors that regulate its production are therefore critical to understand—and potentially even control—the behavior of these bacterial populations.

Within the *S. aureus* species, there are four different *agr* specificity groups that each produce a different AIP molecule (*SI Appendix*, Fig. S1*A*) (5, 11, 12). Although they differ in peptide sequence, each AIP features a conserved thiolactone moiety which is crucial for *agr* activation. The biosynthesis of AIP starts with the translation of AgrD (*SI Appendix*, Fig. S1*B*), which is anchored in the inner leaflet of the cell membrane due to an amphipathic helix located within its N-terminal domain (13). AgrB catalyzes the proteolytic macrothiolactonization of AgrD to produce the key biosynthetic intermediate known as AgrD (1-32) thiolactone (*SI Appendix*, Fig. S2) (14). This reaction is hypothesized to be driven forward thermodynamically by the Clp protease-mediated degradation of the released C-terminal domain (14). In *agr* groups-I, -II, and -IV, the integral membrane protease MroQ cleaves AgrD (1-32) thiolactone to generate the mature AIP pheromone and the N-AgrD leader peptide (*SI Appendix*, Fig. S2), however, the membrane topology of the MroQ active site is unknown (15, 16). Furthermore, the protease that catalyzes the final step of AIP-III production has not been characterized, and the mechanism(s) by which AIP and N-AgrD are secreted out of the cell remains to be discovered.

After the AIP is produced and released, it diffuses until it encounters its cognate AgrC receptor, which subsequently phosphorylates and dimerizes the AgrA transcription factor via a two-component phosphorelay signaling pathway (17, 18). While dimeric AgrA activates the P2 promoter at basal levels, sufficient accumulation of phosphorylated AgrA dimers activates the divergent P3 promoter, resulting in the transcription of a multifunctional RNA called RNAIII (*SI Appendix*, Fig. S3) (19). In addition to encoding δ-toxin, this highly structured RNA is itself also an important regulator of gene expression. For instance, by directly binding to some mRNA targets, RNAIII can block the ribosomal binding site and initiate transcript degradation, leading to gene repression (20). For other transcripts, RNAIII can promote gene expression by either stabilizing the mRNA or by exposing their ribosomal binding sites to increase translation (21, 22).

Activation of AgrA also increases the expression of virulence factors known as phenol-soluble modulins (PSMs) (23). PSMs are an important class of short, amphipathic, α-helical peptides that act as toxins, instigating inflammation and cell lysis within the host. Notably, the N-terminal amphipathic helix of AgrD (1-32) thiolactone resembles a PSM in structure, and indeed, such a classification has been proposed for the N-AgrD leader peptide produced during the biosynthesis of the AIP (24). PSMs are transported out of the cell by the action of two promiscuous ATP-binding cassette (ABC) transporters known as AbcA and PmtCD (*SI Appendix*, Fig. S3) (25– 27). It has been previously observed that mutations in *pmtC* or *abcA* resulted in a loss of hemolytic capacity, which is a hallmark of QS disruption (28). However, a direct biochemical role for these transporters in the *S. aureus* QS system was not investigated (28).

Here, we focus on a critical question that remains unresolved in AIP biosynthesis: how does *S. aureus* secrete the AIP molecule? While various models for AIP release may be postulated (Fig. 1), no experimental result has demonstrated any mechanism to be operative. In this manuscript, we provide evidence that in *agr* groups-I and -II, AbcA represents the major transporter that actively secretes AgrD (1-32) thiolactone out of the cell (in *agr* group-I, PmtCD also exports AgrD-I (1-32) thiolactone, albeit to a more minor extent). Spike-in assays utilizing synthetic AgrD-I (1-32) thiolactone and reconstituted MroQ nanodiscs demonstrate that the active site of MroQ resides within the outer leaflet of the cell membrane, allowing AIPs to be produced on the cell surface directly in the extracellular space. Thus, we propose a new model in which the AbcA/PmtCD transporters directly contribute to *S. aureus* QS by secreting the biosynthetic precursors of certain AIPs into the extracellular milieu where MroQ can catalyze the ultimate proteolysis step.

**Figure 1.**
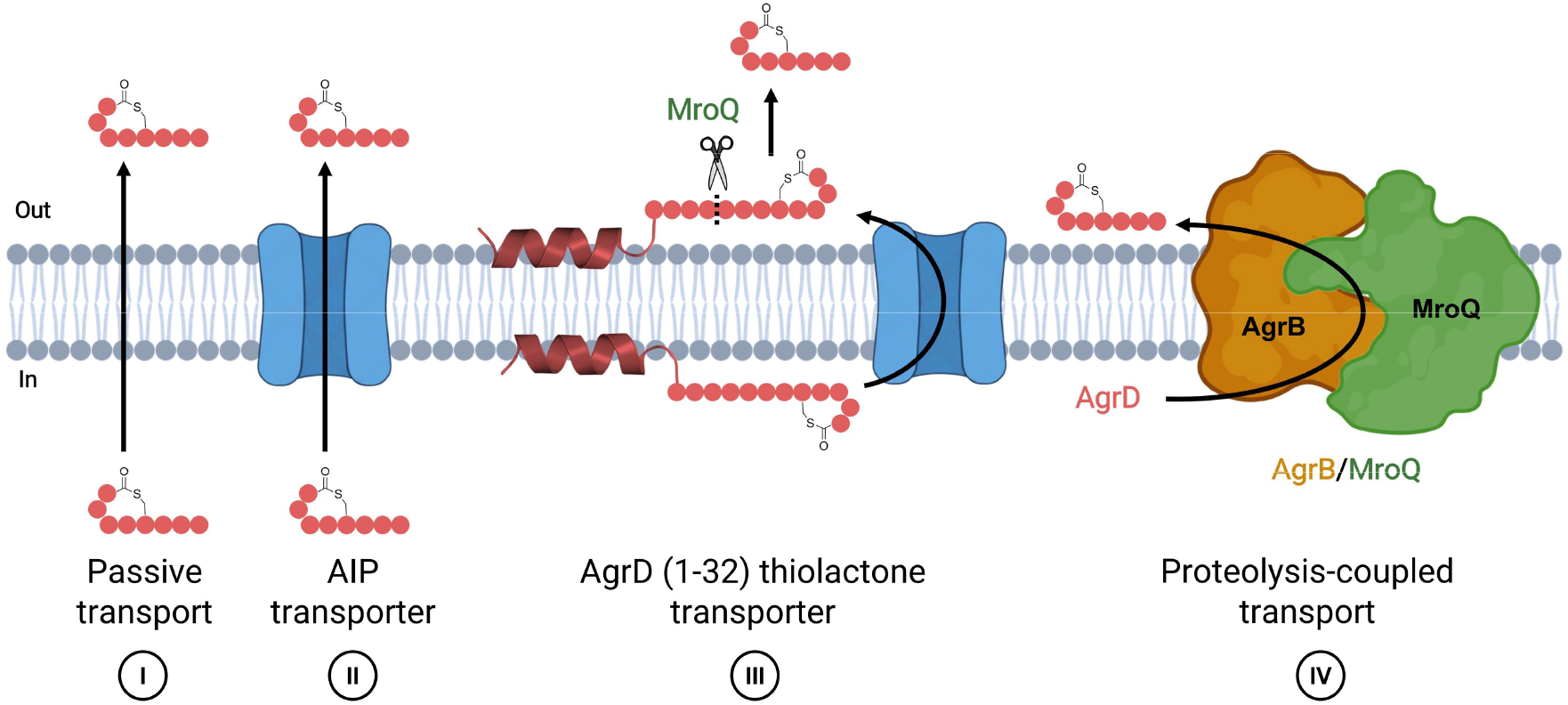
AIP transport models. Four potential models are depicted: (I) Passive diffusion of the mature AIP produced inside the cell through the processing of the precursor peptide AgrD by the integral membrane proteases AgrB and MroQ, (II) Transporter-mediated export of the mature AIP produced inside the cell, (III) Transport of the biosynthetic intermediate AgrD (1-32) thiolactone and its subsequent processing into AIP by the membrane protease MroQ, (IV) Coupled processing and secretion of AgrD by AgrB and MroQ.

## Results

### High-throughput Screening of a Transposon Mutant Library Reveals Regulators of AIP-I production

To identify potential genes that are important for AIP secretion, we took advantage of the Nebraska Transposon (Tn) Mutant Library (NTML), a collection of 1,920 different Tn insertion mutants derived from the *agr* group-I MRSA strain USA300 LAC JE2 (29). First, we optimized and validated a high-throughput assay using a reporter cell strain of *S. aureus* that had been engineered to sensitively detect AIP-I via a colorimetric readout (Fig. 2*A* and *SI Appendix*, Fig. S4) (30). The *agrB* Tn mutant (strain NE95) is unable to produce AIP-I, allowing us to use the basal reporter cell assay signal produced from the conditioned media of strain NE95 to benchmark the signals from all the other strains in the AIP deficiency screen (see *Materials and Methods* for more details). In a similar fashion, the signal produced from the conditioned media of the wild-type (WT) JE2 strain was used as a benchmark in the AIP abundance screen. After subjecting the NTML to our high-throughput reporter cell assay, we assigned an AIP deficiency and abundance score to each Tn mutant (note that higher AIP deficiency scores are associated with lower AIP-I production, whereas higher AIP abundance scores are associated with higher AIP-I production) (Fig. 2*B–C*).

**Figure 2.**
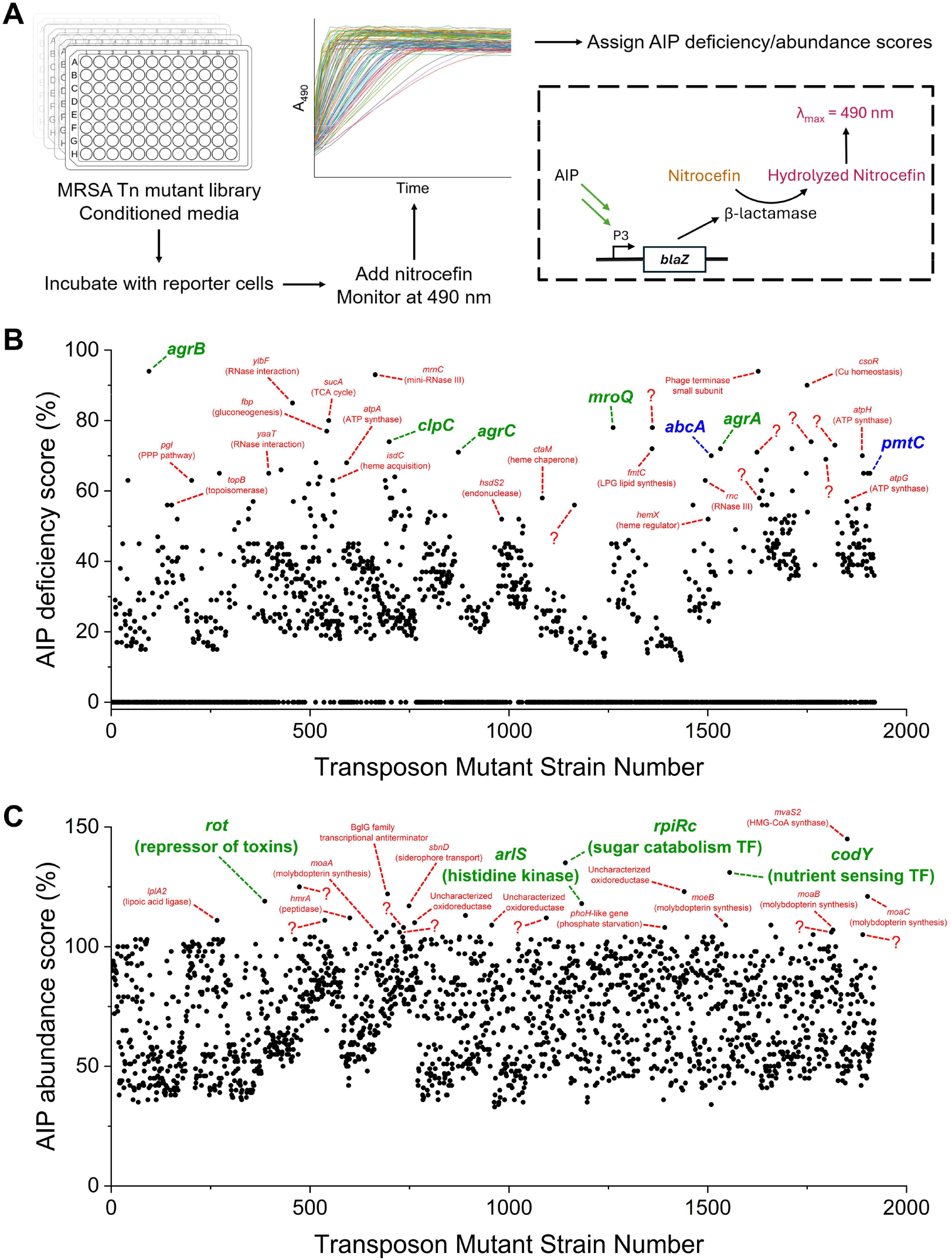
A high-throughput screen for genes involved in AIP production. (*A)* Workflow depicting screening of the Nebraska Transposon Mutant Library (NTML) using a reporter cell assay. The diagram within the dotted box depicts how the *S. aureus* reporter cell allows for the indirect colorimetric detection of AIP-I. (*B* and *C*) High-throughput screening of an *S. aureus* transposon mutant library reveals genes that are important for upregulating (*B*) and downregulating (*C*) AIP-I production. In (*B*), saturated reporter cell signals were assigned an AIP deficiency score of zero. Data is presented as the mean of *n* = 1–4 biological replicates (the assay was repeated for strains with high AIP deficiency/abundance scores for verification purposes).

Gratifyingly, strains with Tn mutations in genes associated with the *agr* system (*agrB, agrC, mroQ*, and *agrA*) had high AIP deficiency scores, as expected. Our screen also identified *clpC* and many ATP synthase subunits as important genes that promote AIP-I production. Tn mutations in genes known to oppose *agr* activation (*rot, arlS, ripRc*, and *codY*) (31–34) resulted in high AIP abundance scores, as did many genes involved in molybdopterin biosynthesis. Many genes not previously known to be related to AIP-I production and several uncharacterized genes were found to be significant hits in our screens. We expect that this dataset will serve as a valuable resource for investigators interested in uncovering novel and previously underappreciated nodes in the regulation of *agr* and staphylococcal virulence (Dataset S1). We were also excited to observe that the two strains with Tn mutations in *abcA* (strain NE1509) and *pmtC* (strain NE1908) had high AIP deficiency scores, suggesting that AbcA and PmtC may be important transporter candidates for AIP-I production.

### AgrD-I (1-32) thiolactone and AgrD-II (1-32) thiolactone are secreted

AbcA and PmtCD are known to actively export PSMs, which share structural similarities with the N-terminal region of AgrD (1-32) thiolactone. Based on this, we hypothesized that these transporters also actively export this biosynthetic intermediate. In this scenario, the processing of AgrD (1-32) thiolactone into AIP and N-AgrD would presumably occur on the exterior cell surface by the action of MroQ, allowing for the release of these two products directly into the extracellular space for *agr* groups-I and -II (third model shown in Fig. 1). To test this hypothesis, we collected cell-free conditioned media from WT and Δ*mroQ* cultures for liquid chromatography-mass spectrometry (LC-MS) analysis, using JE2 as an *agr* group-I model strain and RN6607 as an *agr* group-II model strain. If MroQ is indeed cleaving its substrate on the exterior cell surface, then we would expect gene deletion of *mroQ* to: 1) ablate AIP production, 2) cause accumulation of AgrD (1-32) thiolactone in the outer leaflet of the cell membrane, and 3) facilitate mass transfer of AgrD (1-32) thiolactone from the outer leaflet into the extracellular aqueous compartment where we may be able to detect and quantify this peptide. As anticipated, Δ*mroQ* cultures did not produce any detectable AIP-I or AIP-II in our LC-MS assay, which importantly employs an internal spike-in standard for quantification (Fig. 3*A* and 3*D*, *SI Appendix*, Fig. S5*A*–*C*, and S5*E*). Furthermore, we observed a clear accumulation of AgrD (1-32) thiolactone in the aqueous conditioned media derived from Δ*mroQ* cultures (Fig. 3*B* and 3*E*, *SI Appendix*, Fig. S5*D* and S5*F*). The AgrD (1-32) thiolactone secreted by *S. aureus* was formylated at the N-terminus and peptide identity was confirmed by LC-MS/MS analysis (Fig. 3*C* and 3*F*). This finding agrees with previous reports in which the secreted N-AgrD leader peptide was determined to be N-terminally formylated (35). Importantly, our ability to detect AgrD (1-32) thiolactone in the supernatant of an *mroQ* knockout strain argues against a direct role for this membrane embedded protease in AIP secretion (fourth model shown in Fig. 1) (36).

**Figure 3.**
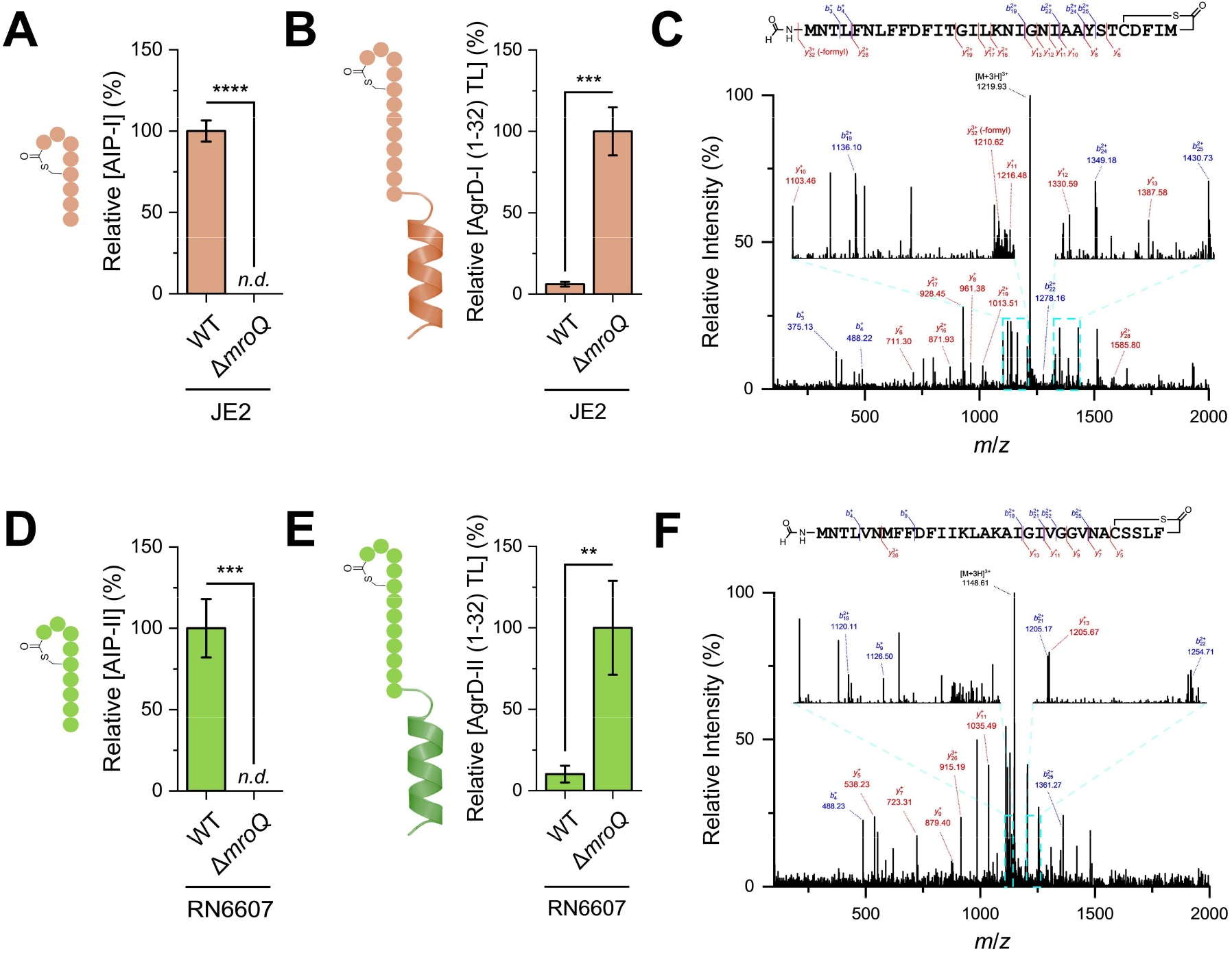
AgrD-I (1-32) thiolactone and AgrD-II (1-32) thiolactone accumulate in the cell-free conditioned media of Δ*mroQ* strains. (*A, B, D*, and *E*) LC-MS quantification of AIP-I (*A*), AgrD-I (1-32) thiolactone (*B*), AIP-II (*D*), and AgrD-II (1-32) thiolactone (*E*) in conditioned media from WT and Δ*mroQ* cultures grown for 18 hrs. Data is presented as mean ± SD (*n* = 3 biological replicates), normalized using both an internal standard and OD_600_, and reported relative to either WT (*A* and *D*) or Δ*mroQ* (*B* and *E*) samples. Statistical analysis is by two-sample *t*-test (****, *P* < 0.0001; ***, *P* < 0.001; **, *P* < 0.01; *n*.*d*. means not detected). (*C* and *F*) Annotated LC-MS/MS fragmentation spectra of Nα-terminally formylated AgrD-I (1-32) thiolactone (*C*) and AgrD-II (1-32) thiolactone (*F*) shown to confirm the identity of the detected peptide species in conditioned media samples from Δ*mroQ* cultures.

### The active site of MroQ faces the extracellular milieu

Next, we sought to establish that the extracellular AgrD-I (1-32) thiolactone we observe can in fact be processed into AIP. As an initial test, we designed an experiment in which MroQ is added exogenously to JE2 Δ*mroQ* cells. In principle, this should lead to rescue of AIP production if the extracellular AgrD-I (1-32) thiolactone is in a structural state that can be processed by the enzyme. To achieve this, we recombinantly expressed and purified WT and catalytically-inactive MroQ fusion proteins (MBP-MroQ-His_6_) from *Escherichia coli* membranes (15) for reconstitution within nanodiscs (discoidal lipid bilayers surrounded by solubilizing membrane scaffolding proteins) (*SI Appendix*, Fig. S6). Consistent with our transport model, AIP-I production was rescued when JE2 Δ*mroQ* cells were incubated with catalytically-active MroQ nanodiscs, but not with the catalytically-inactive mutant version of the protein, demonstrating that extracellular AgrD-I (1-32) thiolactone is the bona fide precursor to AIP-I (Fig. 4*A–C*) (37).

**Figure 4.**
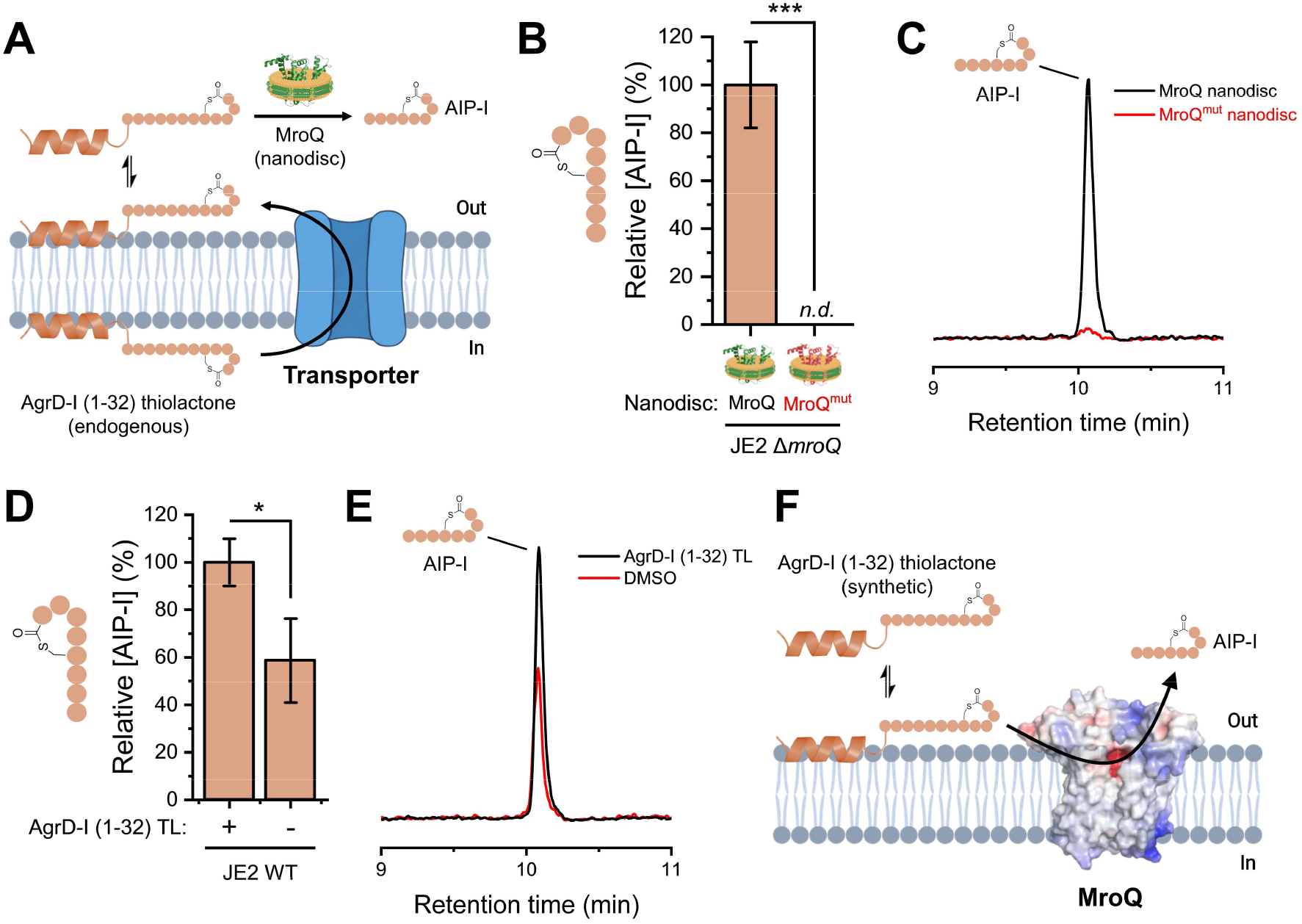
MroQ has an active site facing the extracellular milieu. (*A*) Schematic of the MroQ nanodisc assay used to assess whether exported AgrD-I (1-32) thiolactone can be processed into AIP-I. (B) AIP-I production by JE2 Δ*mroQ* cells grown in media supplemented with either MroQ or MroQ_mut_ nanodiscs. MroQ_mut_ refers to the inactive E141A/E142A double mutant. (*C*) Representative normalized extracted ion chromatogram (EIC) traces for AIP-I used for quantification in (*B*). (*D*) AIP-I production by JE2 WT cells grown in media containing a spike-in of either synthetic AgrD-I (1-32) thiolactone (+) or DMSO (-). (*E*) Representative normalized EIC traces of AIP-I used for quantification in (*D*). (*F*) An electrostatic surface potential model of MroQ cleaving its substrate in the outer leaflet of the cell membrane (key catalytic residues are centered in the middle of the MroQ structure and highlighted as red). In (*B*) and (*D*), data is presented as mean ± SD (*n* = 3 biological replicates), normalized using an internal standard, and reported relative to samples treated with either MroQ nanodiscs (*B*) or synthetic AgrD-I (1-32) thiolactone (*D*). Statistical analysis is by two-sample *t*-test (***, *P* < 0.001; *, *P* < 0.05; *n*.*d*. means not detected). In (*C*) and (*E*), EIC trace intensities are normalized relative to an internal standard. All MroQ structures shown are AlphaFold 3 models (37).

Implicit in our working model is that the active site of MroQ faces the outside of the cell where it can cleave extracellular AgrD-I (1-32) thiolactone. While the MroQ nanodisc experiment shows that extracellular AgrD-I (1-32) thiolactone can be converted into AIP-I, it does not address the question of active site accessibility in the endogenous enzyme. To explore this, we chemically synthesized AgrD-I (1-32) thiolactone by solid-phase peptide synthesis (SPPS) (*SI Appendix*, Fig. S7). The purified peptide was then added to JE2 cultures and the amount of AIP present in the supernatant was measured by LC-MS. We observed increased AIP levels above endogenous levels upon spike-in of the synthetic peptide (Fig. 4*D–E*). The increase in AIP-I production can be attributed to MroQ catalysis because incubating synthetic AgrD-I (1-32) thiolactone with JE2 Δ*mroQ* cells resulted in no detectable AIP-I (*SI Appendix*, Fig. S8). Assuming negligible import of the synthetic peptide, these data indicate that the MroQ active site is accessible to extracellular substrates (Fig. 4*F*).

### AbcA actively exports AgrD-I (1-32) thiolactone and AgrD-II (1-32) thiolactone

The transposon mutant screen suggested that the AbcA and PmtCD transporters play a role in *S. aureus* QS (Fig. 2*B*). To firmly establish the involvement of AbcA and PmtCD in the biosynthesis of certain AIPs,we carried out targeted genetic knockout and rescue experiments. In JE2 cells, gene deletion of these transporters reduced AIP-I levels as measured by LC-MS (*SI Appendix*, Fig. S9). Complementation with ATPase-active versions of these transporters resulted in rescue of AIP-I production, whereas vectors expressing ATPase-inactive transporters (25, 38) did not have the same effect (Fig. 5*A–B* and *SI Appendix*, Fig. S10). When *abcA* and/or *pmtCD* was deleted in a JE2 Δ*mroQ* background, AgrD-I (1-32) thiolactone secretion decreased, with gene deletion of *abcA* resulting in the largest reduction in AgrD-I (1-32) thiolactone levels (*SI Appendix*, Fig. S11*A*–*B*). Gratifyingly, AgrD-I (1-32) thiolactone secretion levels were rescued upon reintroduction of the transporters (Fig. 5*C–D*, *SI Appendix*, Fig. S11*C*–*D*). This data suggests that AbcA and PmtCD both actively export AgrD-I (1-32) thiolactone, with AbcA being the major transporter involved. Intriguingly, when the analogous experiments were performed in RN6607, only AbcA had a significant impact on AIP-II production (Fig. 5*E–F*, *SI Appendix*, Fig. S12) and AgrD-II (1-32) thiolactone secretion (Fig. 5*G–H*, *SI Appendix*, Fig. S13). Thus, we conclude that AbcA is the major exporter of AgrD-II (1-32) thiolactone. AbcA and PmtCD have been reported to have differences in substrate selectivity (25), consistent with our findings of *agr* group-dependent transport mechanisms.

**Figure 5.**
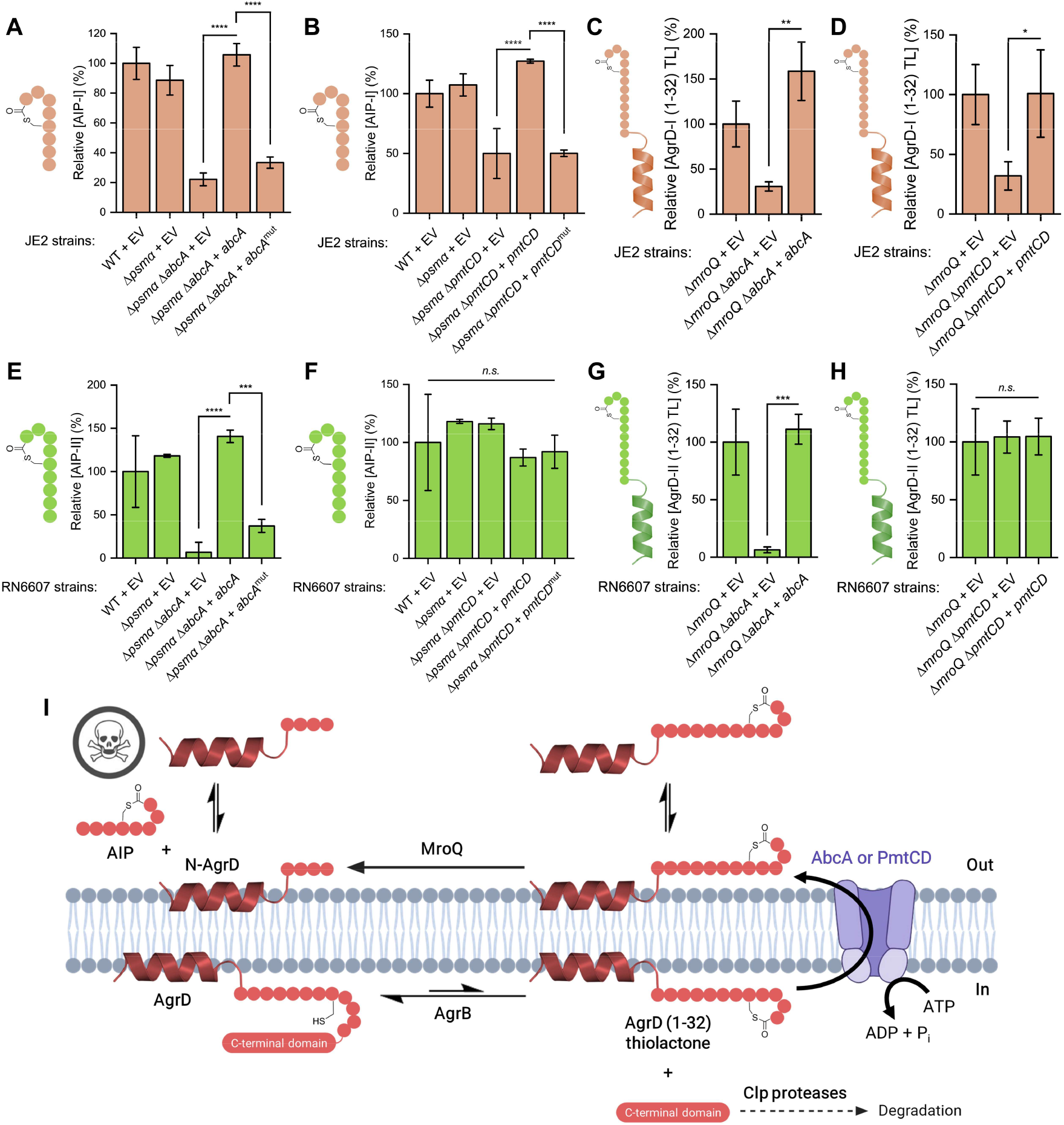
AbcA actively exports the direct precursors of AIP-I and AIP-II. (*A*–*H*) Quantification of AIP and AgrD (1-32) thiolactone in the conditioned media derived from either JE2 (*agr* group-I, panels *A*–*D*) or RN6607 (*agr* group-II, panels *E*–*H*) strains harboring complementation plasmids. Δ*pmtCD* strains had to be generated in either Δ*psmα* or Δ*mroQ* backgrounds to prevent toxic accumulation of membranolytic peptides. EV refers to the empty vector control. Transporters with inactive ATPase domains are referred to as either *abcA*_mut_ (K374M mutant) or *pmtCD*_mut_ (E145Q mutant in PmtC). In (*A*), (*C*), and (*E*–*H*), the complementation plasmids utilize the native *abcA* promoter, whereas in panels (*B*) and (*D*), the complementation plasmids utilize a cadmium-inducible promoter. Cultures were grown for either 12 hrs (*A, C*, and *D*) or 18 hrs (*B* and *E*–*H*). Data is presented as mean ± SD (*n* = 3 biological replicates), normalized using both an internal standard and OD_600_, and reported relative to either WT (*A, B, E*, and *F*) or Δ*mroQ* (*C, D, G*, and *H*) samples. Statistical analysis is by 1-way ANOVA and Tukey HSD post-hoc analysis (****, *P* < 0.0001; ***, *P* < 0.001; **, *P* < 0.01; *, *P* < 0.05; *n*.*s*. means not significant). (*I*) Proposed biosynthetic model for AIP-I, AIP-II, and putatively, AIP-IV.

Gene deletion of *abcA* and/or *pmtCD* in MW2 (an *agr* group-III strain) did not have a large effect on AIP-III production levels (*SI Appendix*, Fig. S14). All attempts to delete *abcA* or *pmtCD* from the genome of RN4850 (an *agr* group-IV strain) were unsuccessful in Δ*psmα* and Δ*mroQ* backgrounds, possibly from the accumulation of toxic PSMβ or δ-toxin peptides (RN4850 is known to have high basal levels of *agr* activation) (12).

## Discussion

It has been known that *S. aureus* secretes AIP to regulate its virulence via QS for thirty years (39), yet the mechanisms detailing how this pheromone is released by cells have remained unknown. The data presented here supports a model in which the biosynthetic intermediate, AgrD (1-32) thiolactone, is actively exported by AbcA in *agr* groups-I and -II, allowing MroQ to catalyze its proteolysis on the cell surface and release AIP and N-AgrD directly into the extracellular milieu (Fig. 5*I*). Our findings also suggest that PmtCD actively exports AgrD-I (1-32) thiolactone, albeit to a lesser extent relative to AbcA.

Gene deletion of *abcA* and/or *pmtCD* does not completely ablate AgrD (1-32) thiolactone secretion, and this may be due to a small amount of cell lysis (resulting in release of intracellular AgrD (1-32) thiolactone), passive diffusion of AgrD (1-32) thiolactone through cell membranes (which remains unexplored), or alternative uncharacterized transport proteins. Although we did not directly interrogate the transport model in *agr* group-IV, we suspect that AIP-IV is released in a similar way to AIP-I given their homology (15). The mechanism by which AIP-III is produced and secreted remains unknown, highlighting the biosynthetic divergence observed in *agr* group-III.

Key to the development of our transport model is the claim that MroQ possesses an active site that resides within the outer leaflet of the cell membrane, allowing it to cleave secreted substrates. In other studies, different membrane topology prediction algorithms gave varying pictures about the location of the MroQ active site, with Phyre2 predicting an active site facing outwards (40) and Protter predicting an active site facing inwards (41). While these bioinformatic analyses were inconclusive, our assays involving spike-in of AgrD-I (1-32) thiolactone and MroQ nanodiscs are more consistent with the active site of MroQ facing outward where it can process the secreted intermediate into the mature AIP. The membrane topology of MroQ also has important implications for its druggability, as anti-virulence inhibitors of MroQ will not have to cross the cell membrane to access the active site.

Some groups have used reporter cell assays to show that *E. coli* expressing *agrBD* can produce AIPs, even though the *E. coli* genome does not have an MroQ homolog (42, 43). Our model resolves this conundrum if AgrD (1-32) thiolactone is secreted by *E. coli* and cleaved by MroQ on reporter cell surfaces, producing AIP *in trans*. In fact, our MroQ nanodisc supplementation assay demonstrates that secreted AgrD-I (1-32) thiolactone can be cleaved to generate AIP-I by the action of MroQ on other membrane surfaces, suggesting that in native *S. aureus* populations, some proportion of exported AgrD (1-32) thiolactone peptides may be processed by neighboring cells (*SI Appendix*, Fig. S15). This type of two-cell metabolic coupling has some precedent (44), and its potential application to *S. aureus* QS yields an interesting perspective on how microbial cells socialize and cooperate.

While our studies add an important missing piece to the *agr* QS puzzle, they also open several interesting questions for future investigations. Do the proteins involved in AIP biosynthesis physically interact within the *S. aureus* cell membrane to improve the efficiency of the overall process? Additional biochemical studies will be needed to investigate this intriguing possibility. Furthermore, much of the AIP-III biosynthetic pathway remains mysterious and represents a significant blind spot for the field that will need to be addressed using genetic and biochemical tools. Finally, beyond the AbcA/PmtCD transporters, our transposon screens revealed many genes that either positively or negatively impact QS. While the involvement of some of these “hits” such as *clpC* can be rationalized based on earlier studies (14), the functional interplay with *agr* QS is far less clear for many others, and many regulatory nodes in the circuit remain undiscovered.

## Materials and Methods

### *S. aureus* Tn mutant screen

Reporter cell assays were performed as previously described (30) with some modifications as detailed in the *SI Appendix*. Briefly, conditioned media from the NTML strains were incubated with RN9222 and nitrocefin, after which the A_490_ was monitored over time. Assay signal is defined as the initial rate of nitrocefin hydrolysis as determined by A_490_. AIP deficiency scores were calculated by dividing the NE95 signal by the signal of the strain of interest (saturated signals were assigned an AIP deficiency score of zero). AIP abundance scores were calculated by dividing the signal of the strain of interest by the WT signal.

### Detection of endogenously produced AIP and AgrD (1-32) thiolactone

Cell pellets from overnight cultures were washed and diluted with sterile phosphate-buffered saline (PBS) such that the OD_600_ of all inocula was 5. 10 µL of inocula was added to 990 µL of media and grown at 37°C, 200 rpm, after which the OD_600_ was measured. Cells were pelleted and the conditioned media was acidified with an internal standard such that the final formic acid concentration was 0.1%. Filtered samples were then submitted for LC-MS analysis.

### MroQ Nanodisc supplementation assay

JE2 Δ*mroQ* cell pellets from overnight cultures were washed and diluted with sterile PBS such that the OD_600_ of all inocula was 5. 0.5 µL of inocula was added to 47 µL of TSB supplemented with 2.5 µL of nanodiscs, after which cultures were grown at 37°C, 200 rpm for 18 hrs. Cells were pelleted, and the conditioned media was acidified with an internal standard such that the final trifluoroacetic acid concentration was 0.1%. Following sample clean-up with solid-phase extraction, the filtered sample was submitted for LC-MS analysis.

### Synthetic AgrD-I (1-32) thiolactone spike-in assay

JE2 cell pellets from overnight cultures were washed and diluted with sterile PBS such that the OD_600_ of all inocula was 5. 0.5 µL of inocula was added to 49 µL of TSB supplemented with 0.5 µL of either synthetic AgrD-I (1-32) thiolactone or DMSO, after which cultures were grown at 37°C, 200 rpm for 5 hrs. Cells were pelleted and the conditioned media was acidified with an internal standard such that the final formic acid concentration was 0.1%. Filtered samples were submitted for LC-MS analysis.

## Data Availability

All study data are included in the article and/or the *SI Appendix*.

## Acknowledgments

We thank current members of the Muir and Gitai laboratories for helpful discussions and comments. We thank Dr. John Eng from the Princeton Chemistry Mass Spectrometry Facility for his assistance. Figures were generated using BioRender and Origin 2026. This work was supported by funds from the Princeton Catalysis Initiative and by a grant from the Novo Nordisk Foundation. A.T. is supported by the National Science Foundation (Graduate Research Fellowship Program Grant No. DGE-2444107) and the Ted Taylor Fellowship (Princeton). K.M.H. is supported by the Center for the Physics of Biological Function (PHY-1734030). Any opinions, findings, and conclusions or recommendations expressed in this material are those of the author(s) and do not necessarily reflect the views of the NSF.

## References

1. M. B. Miller, B. L. Bassler, Quorum sensing in bacteria. Annu. Rev. Microbiol. 55, 165–199 (2001).

2. M. Vestergaard, D. Frees, H. Ingmer, Antibiotic resistance and the MRSA problem. Microbiol. Spectr. 7 (2019).

3. R. P. Novick, E. Geisinger, Quorum sensing in staphylococci. Annu. Rev. Genet. 42, 541–564 (2008).

4. S. P. Bodine, T. W. Muir, Molecular mechanisms of virulence regulation in Staphylococcus aureus: a journey into reconstitutive biochemistry. Acc. Chem. Res. 58, 1657–1669 (2025).

5. P. Mayville et al., Structure-activity analysis of synthetic autoinducing thiolactone peptides from Staphylococcus aureus responsible for virulence. Proc. Natl. Acad. Sci. U.S.A. 96, 1218–1223 (1999).

6. J. M. Yarwood, D. J. Bartels, E. M. Volper, E. P. Greenberg, Quorum sensing in Staphylococcus aureus biofilms. J. Bacteriol. 186, 1838–1850 (2004).

7. A. Björklind, S. Arvidson, Mutants of Staphylococcus aureus affected in the regulation of exoprotein synthesis. FEMS Microbiol. Lett. 7, 203–206 (1980).

8. C. Lebeau, F. Vandenesch, T. Greenland, R. P. Novick, J. Etienne, Coagulase expression in Staphylococcus aureus is positively and negatively modulated by an agr-dependent mechanism. J. Bacteriol. 176, 5534–5536 (1994).

9. B. R. Boles, A. R. Horswill, Agr-mediated dispersal of Staphylococcus aureus biofilms. PLoS Pathog. 4 (2008).

10. P. Recsei et al., Regulation of exoprotein gene expression in Staphylococcus aureus by agr. Mol. Gen. Genet. 202, 58–61 (1986).

11. G. J. Lyon, J. S. Wright, T. W. Muir, R. P. Novick, Key determinants of receptor activation in the agr autoinducing peptides of Staphylococcus aureus. Biochemistry 41, 10095–10104 (2002).

12. S. Jarraud et al., Exfoliatin-producing strains define a fourth agr specificity group in Staphylococcus aureus. J. Bacteriol. 182 (2000).

13. L. Zhang, J. Lin, G. Ji, Membrane anchoring of the AgrD N-terminal amphipathic region is required for its processing to produce a quorum-sensing pheromone in Staphylococcus aureus. J. Biol. Chem. 279, 19448–19456 (2004).

14. B. Wang, A. Zhao, R. P. Novick, T. W. Muir, Key driving forces in the biosynthesis of autoinducing peptides required for staphylococcal virulence. Proc. Natl. Acad. Sci. U.S.A. 112, 10679–10684 (2015).

15. A. Zhao et al., Reconstitution of the S. aureus agr quorum sensing pathway reveals a direct role for the integral membrane protease MroQ in pheromone biosynthesis. Proc. Natl. Acad. Sci. U.S.A. 119 (2022).

16. M. R. Stock et al., Characterization of MroQ-dependent maturation and export of the Staphylococcus aureus accessory gene regulatory system autoinducing peptide. Infect. Immun. 90 (2022).

17. R. P. Novick et al., The agr P2 operon: an autocatalytic sensory transduction system in Staphylococcus aureus. Mol. Gen. Genet. 248, 446–458 (1995).

18. K. Rajasree, A. Fasim, B. Gopal, Conformational features of the Staphylococcus aureus AgrA-promoter interactions rationalize quorum-sensing triggered gene expression. Biochem. Biophys. Rep. 6, 124–134 (2016).

19. R. P. Novick et al., Synthesis of staphylococcal virulence factors is controlled by a regulatory RNA molecule. EMBO J. 12, 3967–3975 (1993).

20. C. Chevalier et al., Staphylococcus aureus RNAIII binds to two distant regions of coa mRNA to arrest translation and promote mRNA degradation. PLoS Pathog. 6 (2010).

21. R. K. Gupta, T. T. Luong, C. Y. Lee, RNAIII of the Staphylococcus aureus agr system activates global regulator MgrA by stabilizing mRNA. Proc. Natl. Acad. Sci. U.S.A. 112, 14036–14041 (2015).

22. E. Morfeldt, D. Taylor, A. von Gabain, S. Arvidson, Activation of alpha-toxin translation in Staphylococcus aureus by the trans-encoded antisense RNA, RNAIII. EMBO J. 14, 4569–4577 (1995).

23. S. Y. Queck et al., RNAIII-independent target gene control by the agr quorum-sensing system: insight into the evolution of virulence regulation in Staphylococcus aureus. Mol. Cell. 32, 150–158 (2008).

24. K. Schwartz et al., The AgrD N-terminal leader peptide of Staphylococcus aureus has cytolytic and amyloidogenic properties. Infect. Immun. 82, 3837–3844 (2014).

25. S. W. Dickey, D. J. Burgin, S. Huang, D. Maguire, M. Otto, Two transporters cooperate to secrete amphipathic peptides from the cytoplasmic and membranous milieus. Proc. Natl. Acad. Sci. U.S.A. 120 (2023).

26. S. S. Chatterjee et al., Essential Staphylococcus aureus toxin export system. Nat. Med. 19, 364–367 (2013).

27. H. Yoshikai et al., Multidrug-resistance transporter AbcA secretes Staphylococcus aureus cytolytic toxins. J. Infect. Dis. 213, 295–304 (2016).

28. Y. Liu et al., Export by a quorum sensing–controlled drug exporter underlies rifampicin’s strong activity against staphylococcal biofilms. Sci. Transl. Med. 17 (2025).

29. P. D. Fey et al., A genetic resource for rapid and comprehensive phenotype screening of nonessential Staphylococcus aureus genes. mBio. 4 (2013).

30. G. J. Lyon, P. Mayville, T. W. Muir, R. P. Novick, Rational design of a global inhibitor of the virulence response in Staphylococcus aureus, based in part on localization of the site of inhibition to the receptor-histidine kinase, AgrC. Proc. Natl. Acad. Sci. U.S.A. 97, 13330–13335 (2000).

31. B. Saïd-Salim et al., Global regulation of Staphylococcus aureus genes by Rot. J. Bacteriol. 185, 610–619 (2003).

32. B. Fournier, A. Klier, G. Rapoport, The two-component system ArlS–ArlR is a regulator of virulence gene expression in Staphylococcus aureus, 41, 247–261 (2001).

33. D. Balasubramanian et al., Staphylococcus aureus coordinates leukocidin expression and pathogenesis by sensing metabolic fluxes via RpiRc. mBio. 7 (2016).

34. C. D. Majerczyk et al., Direct targets of CodY in Staphylococcus aureus. J Bacteriol. 192, 2861–2877 (2010).

35. D. J. Gonzalez et al., N-terminal ArgD peptides from the classical Staphylococcus aureus agr system have cytotoxic and proinflammatory activities. 21, 1457–1462 (2014).

36. L. Fang, C. Cosgriff, F. Alonzo, 3rd, Determinants of maturation of the Staphylococcus aureus autoinducing peptide. J. Bacteriol. 206 (2024).

37. J. Abramson et al., Accurate structure prediction of biomolecular interactions with AlphaFold 3. Nature. 630 493–500 (2024).

38. N. Zeytuni et al., Structural insight into the Staphylococcus aureus ATP-driven exporter of virulent peptide toxins. Sci. Adv. 6 (2020).

39. G. Ji, R. C. Beavis, R. P. Novick, Cell density control of staphylococcal virulence mediated by an octapeptide pheromone. Proc. Natl. Acad. Sci. U.S.A. 92, 12055–12059 (1995).

40. C. J. Cosgriff, C. R. White, W. P. Teoh, J. P. Grayczyk, F. Alonzo, 3rd, Control of Staphylococcus aureus quorum sensing by a membrane-embedded peptidase. Infect. Immun. 87 (2019).

41. S. Marroquin et al., MroQ is a novel Abi-domain protein that influences virulence gene expression in Staphylococcus aureus via modulation of agr activity. Infect. Immun. 87 (2019).

42. M. Thoendel, A. R. Horswill, Identification of Staphylococcus aureus AgrD residues required for autoinducing peptide biosynthesis. J. Biol. Chem. 284, 21828–21838 (2009).

43. N. Marchand, C. H. Collins, Peptide-based communication system enables Escherichia coli to Bacillus megaterium interspecies signaling. Biotechnol. Bioeng. 110, 3003–3012 (2013).

44. S. Zhang et al., A chemical radar allows bacteria to detect and kill predators. Cell. 188, 2495–2504 (2025).

