## Supplementary material for "Release of the *Staphylococcus aureus* autoinducing peptide is enabled by active export of the biosynthetic intermediate": SI Appendix

### Supporting Information Text

### Extended Methods

***Materials and General Methods.*** All common buffers and salts, Tris(2-carboxyethyl)phosphine hydrochloride (TCEP), phenylmethylsulfonyl fluoride (PMSF), isopropyl-β-D-thiogalactopyranoside (IPTG), Luria-Bertani (LB) broth, LB agar, Coomassie brilliant blue (CBB), yeast extract, nitrocefin, SilverQuest silver staining kit, and HisPur Nickel-nitrilotriacetic acid (Ni-NTA) resin were purchased from ThermoFisher Scientific (Waltham, MA). Ampicillin and kanamycin were purchased from GoldBio (St. Louis, MO). Tryptic soy agar (TSA), 1,2-dimyristoyl-*sn*-glycero-3-phosphocholine (DMPC), and 1,2-dimyristoyl-sn-glycero-3-phosphoglycerol (DMPG) were purchased from VWR (Radnor, PA). Tryptic soy broth (TSB) was purchased from Genesee Scientific (El Cajon, CA). Casamino acids were purchased from Difco Biosciences, Inc. (San Diego, CA). β-glycerophosphate was purchased from EMD Millipore (Billerica, MA). Trifluoroacetic acid (TFA) was purchased from Halocarbon (North Augusta, SC). Ethyl 2-cyano-2-(hydroxyimino)acetate (Oxyma), 5-bromo-4-chloro-3-indolyl-β-D-galactopyranoside (X-Gal), tetradecylphosphocholine (FC-14) and dodecylphosphocholine (FC-12) were purchased from Ambeed (Buffalo Grove, IL). Glycerol was purchased from Neta Scientific (Marlton, NJ). Fmoc-protected amino acids were purchased from Matrix Innovation (Quebec, Canada). Boc-L-methionine was purchased from Novabiochem (Nottingham, UK). *N*,*N′*-diisopropylcarbodiimide (DIC) was purchased from Chem-Impex (Wood Dale, IL). Oligonucleotides were purchased from either Integrated DNA Technologies (Coralville, IA) or MilliporeSigma (Burlington, MA). Gibson Assembly Master Mix and all empty plasmids were purchased from New England Biolabs (Ipswich, MA). Stbl3, DH5α, BL21(DE3), and C43(DE3) *Escherichia coli* strains were purchased from Invitrogen (Carlsbad, CA) and subcultured to generate stocks of competent cells. BL21(DE3)-RIL cells used for TEV protease expression were acquired from Addgene (Watertown, MA). The *S. aureus* transposon library (Nebraska Transposon Mutant Library (NTML) Screening Array, NR-48501) and RN4850 (NR-45955) were obtained from the Network on Antimicrobial Resistance in *Staphylococcus aureus* (NARSA) for distribution through BEI Resources, NIAID, NIH (Manassas, VA). Ammonium persulfate (APS), 30% Acrylamide/Bis Solution 29:1, tetramethylethylenediamine (TEMED), criterion empty gel cassettes, and Bio-Beads SM2 were purchased from Bio-Rad (Hercules, CA). Sep-Pak $C_{18}$ vac cartridges (1 cc, 100 mg, 55-105 μm) were purchased from Waters (Milford, MA). Amicon MWCO centrifugal filter unit concentrators were obtained from MilliporeSigma (Burlington, MA). Empty fritted chromatography columns were purchased from Environmental Express (Charleston, SC). DNA purification kits were purchased from Qiagen (Valencia, CA). All other reagents and solvents were purchased from Sigma-Aldrich (St. Louis, MO). All reagents and solvents were used without further purification. Plasmid sequences and all *S. aureus* strains used in this study were validated by sequencing performed by either Genewiz (South Plainfield, NJ) or Plasmidsaurus (South San Francisco, CA).

Cell disruption was performed using an ATA Scientific Instruments Avestin Emulsiflex C3 French Press (Caringbach NSW, Australia). Size-exclusion chromatography (SEC) was performed with an ÄKTA FPLC equipped with a Frac-950 fraction collector using either a Superdex S200 10/300 GL or a Superose 6 Increase 10/300 GL column, and anion exchange chromatography was performed using a Hiprep Q FF 16/10 column from Cytiva (Marlborough, MA). Sodium dodecyl sulfate-polyacrylamide gel electrophoresis (SDS-PAGE) gels were imaged on a Cytiva Amersham ImageQuant 800 (Malborough, MA). For solid-phase peptide synthesis (SPPS) of AgrD-I (1-32) thiolactone, a Liberty Blue Peptide Synthesizer from CEM was used (Matthews, NC). Fmoc-SPPS of the leu-enkephalin (H-YGGFL-$NH_2$) internal standard was performed manually following standard procedures. Large scale peptide purification was performed by preparative-scale reverse-phase high-performance liquid chromatography (RP-HPLC) using a 2535 Quartenary Gradient Module equipped with a 2489 UV/Vis Detector and a Waters Fraction Collector III employing an XBridge peptide bridged ethylene hybrid (BEH) $C_{18}$ optimum bed density (OBD) preparative column (300Å,10 μm, 19 × 250 mm) from Waters (Milford, MA). Large-scale purification was performed using 90% water + 10% isopropyl alcohol (IPA) + 0.1% TFA as solvent A and 90% acetonitrile + 10% IPA + 0.1% TFA as solvent B. Synthetic peptides were analyzed at small scale utilizing analytical-scale RP-HPLC on an Agilent 1260 Infinity HPLC system employing an Eclipse Plus $C_{18}$

column (5 µm, 4 × 150 mm) from Agilent (Santa Clara, CA). Small-scale RP-HPLC was performed using water + 0.1% TFA as solvent A and 90% acetonitrile + 10% water + 0.1% TFA as solvent B. Mass validation of synthetic peptides was performed by electrospray ionization mass spectrometry (ESI-MS) using a 6120 Quadrupole liquid-chromatography-mass spectrometry (LC-MS) employing a 1260 Infinity LC system with a Zorbax 300SB-$C_{18}$ column (3.5 µm, 4.6 × 100 mm) from Agilent (Santa Clara, CA). Conditioned media samples were analyzed by LC-MS(/MS) using an Agilent 6546 LC/Q-TOF employing a 1290 Infinity II series system running a gradient elution method with mobile phases consisting of water + 0.1% formic acid (FA) as solvent A and acetonitrile + 0.1% FA as solvent B. For LC-MS analyses, the column utilized was either a Vydac 214TP $C_4$ column (5 µm, 150 × 4.6 mm) or a YMC-Pack phenyl column (5 µm, 50 × 4.6 mm). Lyophilization of samples after preparative RP-HPLC or solid phase extraction (SPE) was performed on a Millrock Technology MD85 lyophilizer (Kingston, NY). Analysis of nitrocefin cleavage in β-lactamase *agr* activation reporter cell assays was performed using either a Biotek microplate reader from Agilent (Santa Clara, CA) or a Tecan microplate reader from Tecan (Männedorf, Switzerland).

***Bacterial strains and plasmids.*** The bacterial strains, plasmids, and primers used in this study are listed in Supplementary Tables S1–3. Unless otherwise noted, *E. coli* strains were routinely grown in LB and LB agar, and *S. aureus* strains were routinely grown in TSB and TSA. When appropriate, media was supplemented with ampicillin (100 µg/mL), kanamycin (50 µg/mL), erythromycin (10 µg/mL), X-Gal (100 µg/mL) or $CdCl_2$ (10 µM). Standard cloning procedures (Gibson assembly, ligation cloning, site-directed mutagenesis, etc.) were performed using either DH5α or Stbl3 cells. Plasmids for *S. aureus* were first electroporated into *S. aureus* RN4220, then phage transduced into the desired host strain using phage 80α. Gene deletion mutants of *S. aureus* were generated via allelic exchange with the pMAD shuttle vector (1) using the double-crossover method. Briefly, pMAD plasmids were cloned to contain the sequences upstream (~1 kb) and downstream (~1 kb) of the gene to be deleted. After phage transduction into the strain of interest (30°C on TSA + erythromycin), a colony was streaked for growth on TSA + erythromycin at 44°C. Then, a colony was grown in TSB (30°C, 200 rpm), after which dilutions were plated on TSA + X-Gal (44°C). White colonies were picked to check for gene deletion by PCR and sequencing, and erythromycin sensitivity was confirmed to ensure proper plasmid curing. Complementation plasmids were generated with either the pCN51 vector containing a cadmium-inducible promoter (2) or a version of the pOS1 vector (3) modified to contain the native JE2 promoter sequence of *abcA*.

**S. aureus *Tn mutant screen*.** Overnight cultures of NTML strains were grown in CYGP media without glucose, diluted 100-fold into fresh media, and grown for 5 hrs with gentle rocking at 37°C, after which cells were pelleted to isolate the conditioned media. The *agr* group-I reporter cell line (RN9222) was grown shaking at 37°C to an $OD_{600}$ of 0.4-0.6. For the AIP deficiency screen, 50 µL of conditioned media was incubated with 50 µL of reporter cells in 96-well plates at 37°C for 1 hr with gentle shaking. For the AIP abundance screen, 5 µL of conditioned media was diluted with 45 µL of media prior to incubation with 50 µL of reporter cells to prevent signal saturation. Then, 100 µL of 100 µM nitrocefin prepared in assay buffer (100 mM phosphate, 5 mM $NaN_3$, 20% propylene glycol, pH 6.0) was added and the $A_{490}$ was monitored over time at 37°C. Assay signal is defined as the initial rate of nitrocefin hydrolysis as determined by $A_{490}$. AIP deficiency scores were calculated by dividing the NE95 signal by the signal of the strain of interest (saturated signals were assigned an AIP deficiency score of zero). AIP abundance scores were calculated by dividing the signal of the strain of interest by the WT signal.

***Detection of endogenously produced AIP and AgrD (1-32) thiolactone.*** Single colonies of the indicated strains were picked and grown in tryptic soy broth (TSB) overnight at 37°C, 200 rpm. Erythromycin (10 µg/mL) was included in the media for strains harboring complementation plasmids and $CdCl_2$ (10 µM) was included for strains harboring plasmids with cadmium-inducible promoters. Cell pellets from the overnight cultures were washed 2-3 times with sterile phosphate-buffered saline (PBS) and diluted such that the $OD_{600}$ of all inocula was 5. 10 µL of inocula was added to 990 µL of media and grown at 37°C, 200 rpm for the indicated times, after which cultures were kept on ice to halt growth prior to measuring $OD_{600}$. Cells were pelleted and 72 µL of conditioned media was acidified with 8 µL of internal standard solution (20-100 nM H-YGGFL-$NH_2$ in 1% FA in water). Samples were filtered (0.45 µm PTFE spin filter) and 20 µL was injected for LC-MS analysis.

***Nanodisc assembly and purification.*** MSP1E3D1(-) was prepared according to previously published protocols (4, 5). Briefly, His-tagged MSP1E3D1 was expressed in BL21(DE3) and purified by Ni-NTA. The His-tag was removed by TEV protease digestion (6), and reverse Ni-NTA purification followed by anion exchange chromatography (Hiprep Q FF 16/10) yielded MSP1E3D1(-). Detergent-solubilized MBP-MroQ-$His_6$ and MBP-MroQ(E141A/E142A)-$His_6$ were prepared as previously described (7). Briefly, MroQ fusion proteins were expressed in C43(DE3) membranes, extracted with detergent, and purified by Ni-NTA and SEC (Superdex S200 10/300 GL). The lipid stock solution consisted of a 1:3 mixture of DMPC and DMPG dissolved in buffer (20 mM HEPES, 100 mM NaCl, 1 mM TCEP, 100 mM sodium cholate, pH 7.5). MSP1E3D1(-), MroQ fusion proteins, and lipids were mixed at a final molar ratio of 4:1:500, ensuring that the final sodium cholate concentration was adjusted to 20 mM. In the pre-assembly mix, MSP1E3D1(-) was incubated with lipids and supplemental sodium cholate, and MroQ fusion proteins were incubated with 0.2% (w/v) FC-12 separately at 30°C for 15 min. The two mixtures were then combined and incubated at 30°C for 30 min. Then, adsorbent SM-2 Bio-Beads (50× the mass of sodium cholate) were added to remove detergent from the system and facilitate spontaneous nanodisc formation (30°C, 2 hr with stirring). Following Bio-Bead elution and Ni-NTA purification to wash away empty nanodiscs, the elution was further purified and characterized using SEC (Superose 6 Increase 10/300 GL) and SDS-PAGE.

***MroQ Nanodisc supplementation assay.*** Single colonies of JE2 Δ*mroQ* were picked and grown in TSB overnight at 37°C, 200 rpm. Cell pellets from the overnight cultures were washed 3 times with sterile PBS and diluted such that the $OD_{600}$ of all inocula was 5. Purified nanodiscs harboring either MBP-MroQ-$His_6$ or MBP-MroQ(E141A/E142A)-$His_6$ were prepared as 2 µM stock solutions in buffer (20 mM Tris, 100 mM NaCl, 1 mM TCEP, 10% glycerol, pH 7.5). 0.5 µL of inocula was added to 47 µL of TSB supplemented with 2.5 µL of nanodiscs, after which cultures were grown at 37°C, 200 rpm for 18 hrs. Cells were pelleted, and 45 µL of conditioned media was acidified with 5 µL of internal standard solution (400 nM H-YGGFL-$NH_2$ in water + 0.1% TFA). Samples were applied to a $C_{18}$ solid-phase extraction (SPE) cartridge pre-conditioned with 4 mL of acetonitrile + 0.1% TFA and 2 mL of water + 0.1% TFA. The SPE cartridge was then washed with 1 mL of water + 0.1% TFA and eluted with 3× 1 mL of acetonitrile + 0.1% TFA. The elution was lyophilized, re-dissolved in 30 µL of water + 0.1% FA, filtered through a 0.45 µm PTFE filter, and 20 µL was injected for LC-MS analysis.

***Synthesis of AgrD-I (1-32) thiolactone.*** Synthetic AgrD-I (1-32) thiolactone was prepared following a modified protocol from previous studies (8, 9) with differences highlighted herein. Fmoc-MeDbz was synthesized from 3-amino-4-(methylamino)-benzoic acid (8), and the MeDbz-Gly-resin was synthesized (9) using the Aminomethyl ChemMatrix resin. Following manual coupling of the C-terminal amino acid (Fmoc-protected methionine), microwave-assisted SPPS allowed for automated peptide elongation using standard DIC/Oxyma chemistry (note that the N-terminal methionine is Boc-protected). Activation of the linker with 4-nitrophenyl chloroformate and *N*,*N*-diisopropylethylamine (DIPEA) followed by global deprotection (TFA/thioanisole/triisopropylsilane (TIPS)/water in a v/v ratio of 89:5:3:3) furnished the peptidyl-*N*-acyl-*N*'-methyl-benzimidazolinone (MeNbz)-Gly-resin. Shaking this resin in a buffered acetonitrile solution (a 1:1 buffer:acetonitrile solution where the buffer consisted of 200 mM sodium phosphate, 2.5% thiodiglycol, pH 6.8) at 50°C for 2 hrs induced macrothiolactonization and peptide release. After resin removal, the crude AgrD-I (1-32) thiolactone was purified by preparative RP-HPLC and characterized by analytical RP-HPLC and ESI-MS.

***Synthetic AgrD-I (1-32) thiolactone spike-in assay.*** Single colonies of either JE2 or JE2 Δ*mroQ* were picked and grown in TSB overnight at 37°C, 200 rpm. Cell pellets from the overnight cultures were washed 3 times with sterile PBS and diluted such that the $OD_{600}$ of all inocula was 5. Purified synthetic AgrD-I (1-32) thiolactone was prepared as a 500 µM stock solution in DMSO. 0.5 µL of inocula was added to 49 µL of TSB supplemented with 0.5 µL of either synthetic AgrD-I (1-32) thiolactone or DMSO, after which cultures were grown at 37°C, 200 rpm for 5 hrs. Cells were pelleted and 36 µL of conditioned media was acidified with 4 µL of internal standard solution (400 nM of H-YGGFL-$NH_2$ in 1% FA). Samples were filtered (0.45 µm PTFE spin filter) and 20 µL was injected for LC-MS analysis.

***LC-MS data processing for analyte quantification.*** Extracted ion chromatograms (EICs) selecting for the mass of each analyte of interest and the internal standard were generated for each analyzed sample and smoothed in the Masshunter Workstation software from Agilent (Santa Clara, CA). Peak areas were obtained by integration, and analyte levels are normalized by dividing the peak area of the analyte of interest by the peak area of the internal standard and/or the $OD_{600}$ of the culture from which the sample was taken. The data is then reported relative to a sample group as specified in the figure legends.

**Figures**

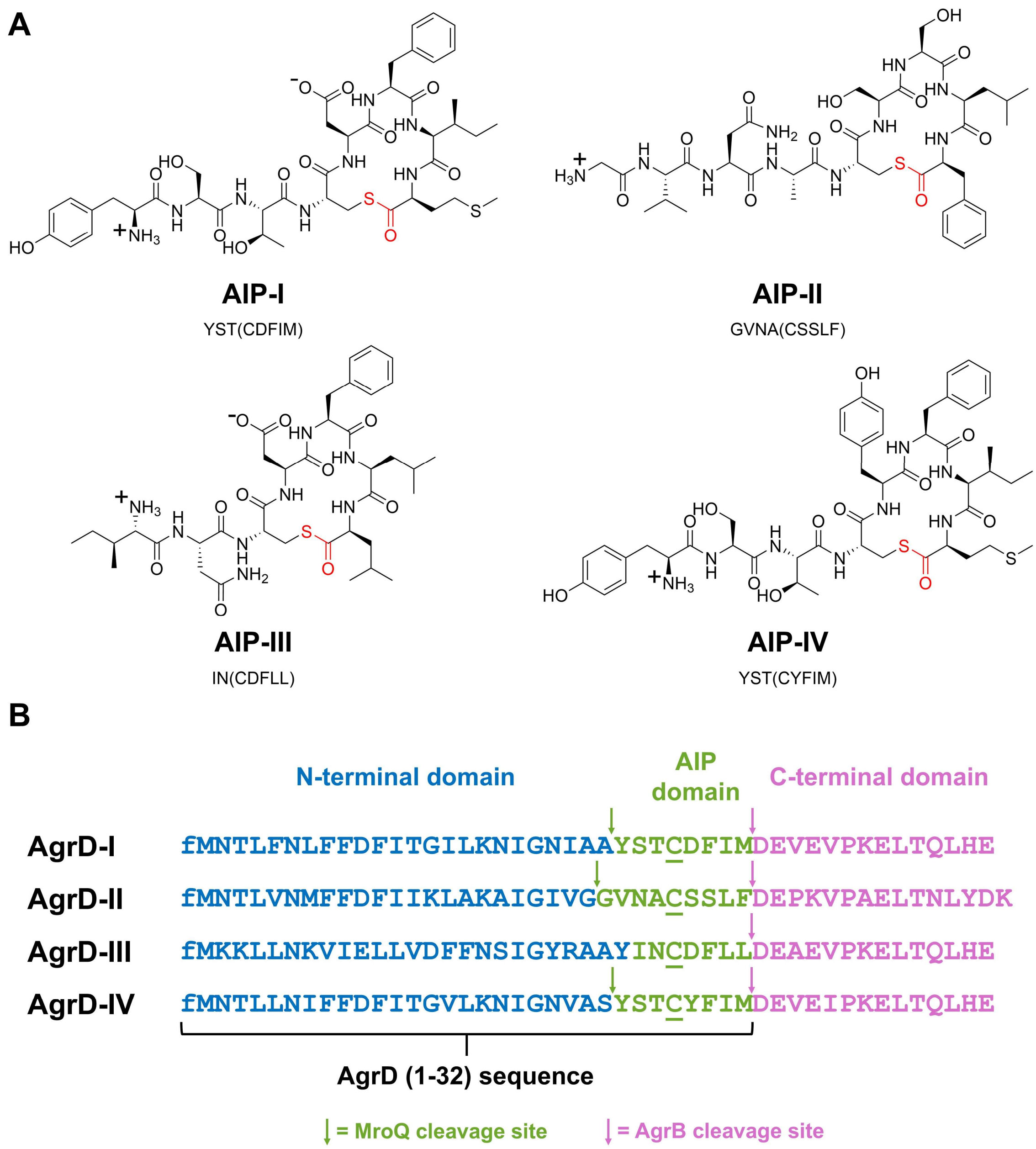


**Figure S1.** The four AIPs and AgrD sequences of *S. aureus*. (*A*) Structures and sequences of the four AIP molecules, highlighting the conserved thiolactone moiety in red. The residues within the macrocycle are shown within parentheses in the sequence. (*B*) The four AgrD sequences, with the key conserved cysteine underlined and the MroQ/AgrB processing sites indicated. UniprotKB accession numbers for AgrD Sequences: AgrD-I (A0A0H2XGM7), AgrD-II (O33586), AgrD-III (O33589), and AgrD-IV (Q9L561).

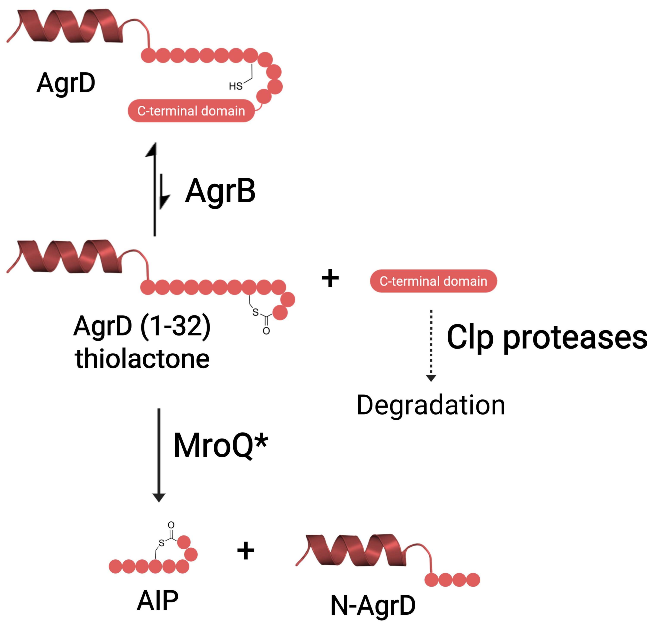


**Figure S2.** Generalized scheme of AIP biosynthesis. *MroQ does not participate in the maturation of AIP-III.

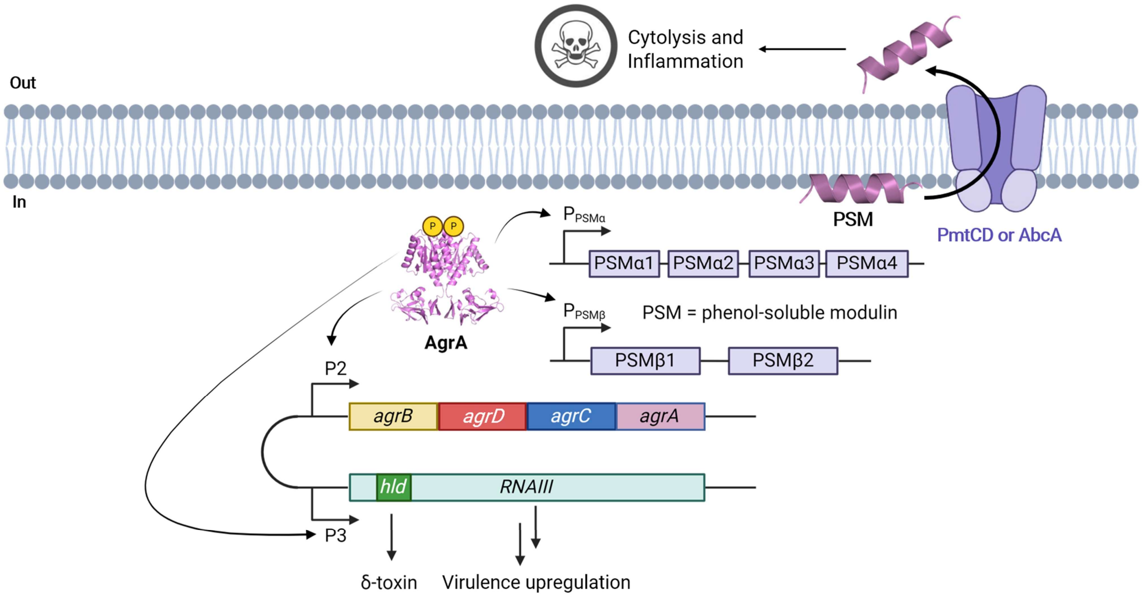


**Figure S3.** AgrA activation upregulates the expression of many genes, resulting in virulent behavior. Notably, the phenol-soluble modulins (PSMs) are actively exported by the AbcA/PmtCD transporters. The AgrA dimer structure shown is an AlphaFold 3 model (10).

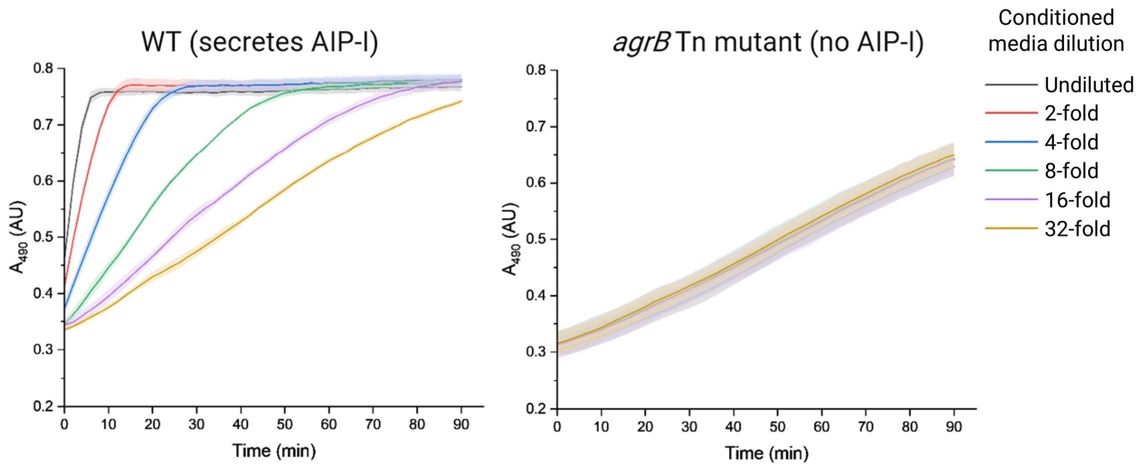


**Figure S4.** Reporter cell validation assay. Conditioned media derived from wild-type (WT) and *agrB* Tn mutant (strain NE95) cultures were diluted as indicated and subjected to the reporter cell assay, validating that this method can robustly distinguish between strains that can produce AIP-I and strains that cannot. Data is presented as mean with error bands representing SD ($n$ = 3 technical replicates).

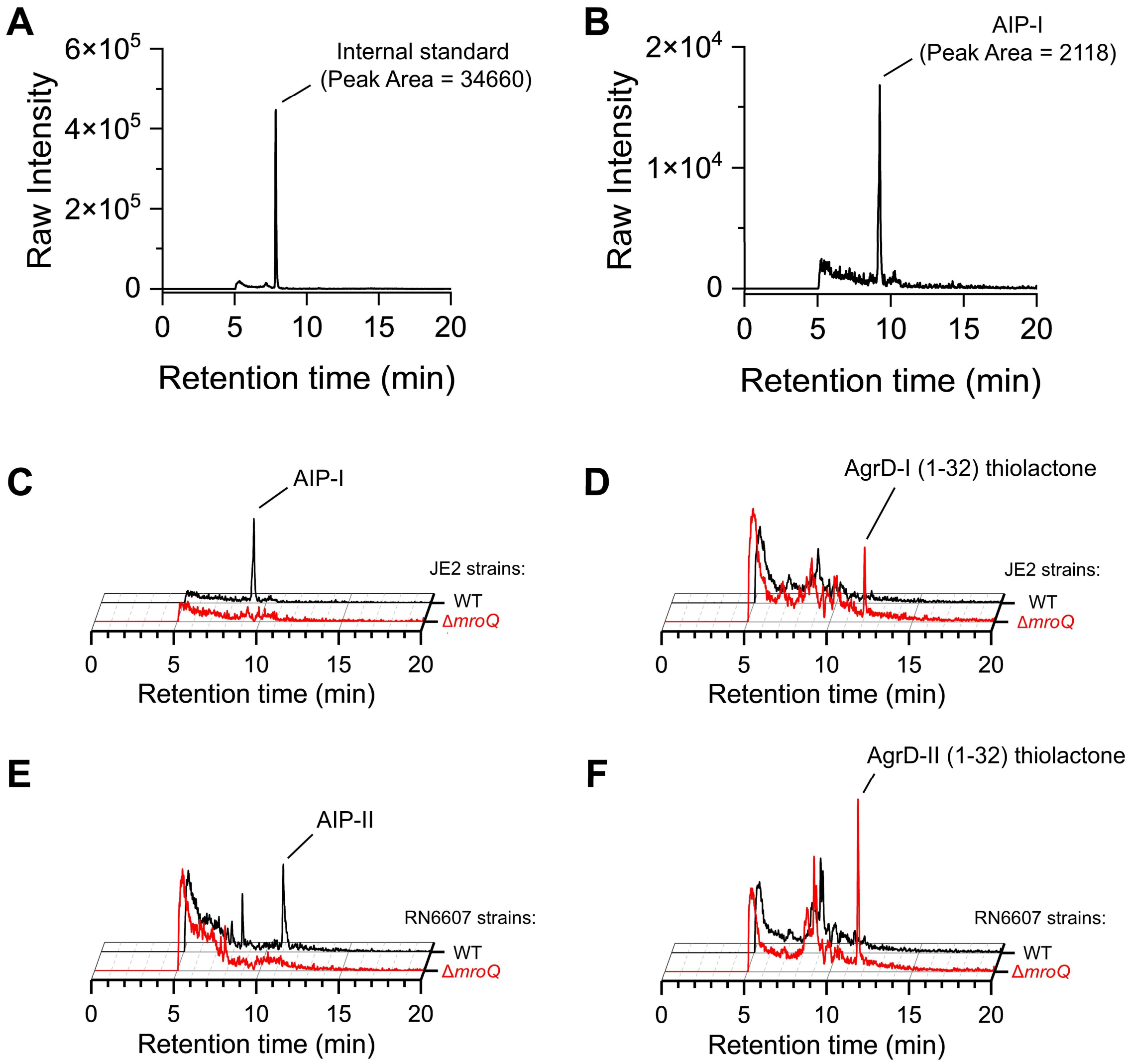


**Figure S5.** Detection and quantification of AIP and AgrD (1-32) thiolactone in conditioned media. (*A* and *B*) Example EIC traces of the leu-enkephalin (H-YGGFL-$NH_2$) internal standard (*A*) and AIP-I (*B*) used for analyte quantification. Peak areas were obtained by integration, and analyte levels shown in this manuscript are normalized by dividing the peak area of the analyte of interest by the peak area of the internal standard and/or the $OD_{600}$ of the culture from which the sample was taken. The data is then reported relative to a sample group as specified in the figure legends. (*C*) Representative normalized EIC traces used for quantification in Fig. 3*A*. (*D*) Representative normalized EIC traces used for quantification in Fig. 3*B*. (*E*) Representative normalized EIC traces used for quantification in Fig. 3*D*. (*F*) Representative normalized EIC traces used for quantification in Fig. 3*E*.

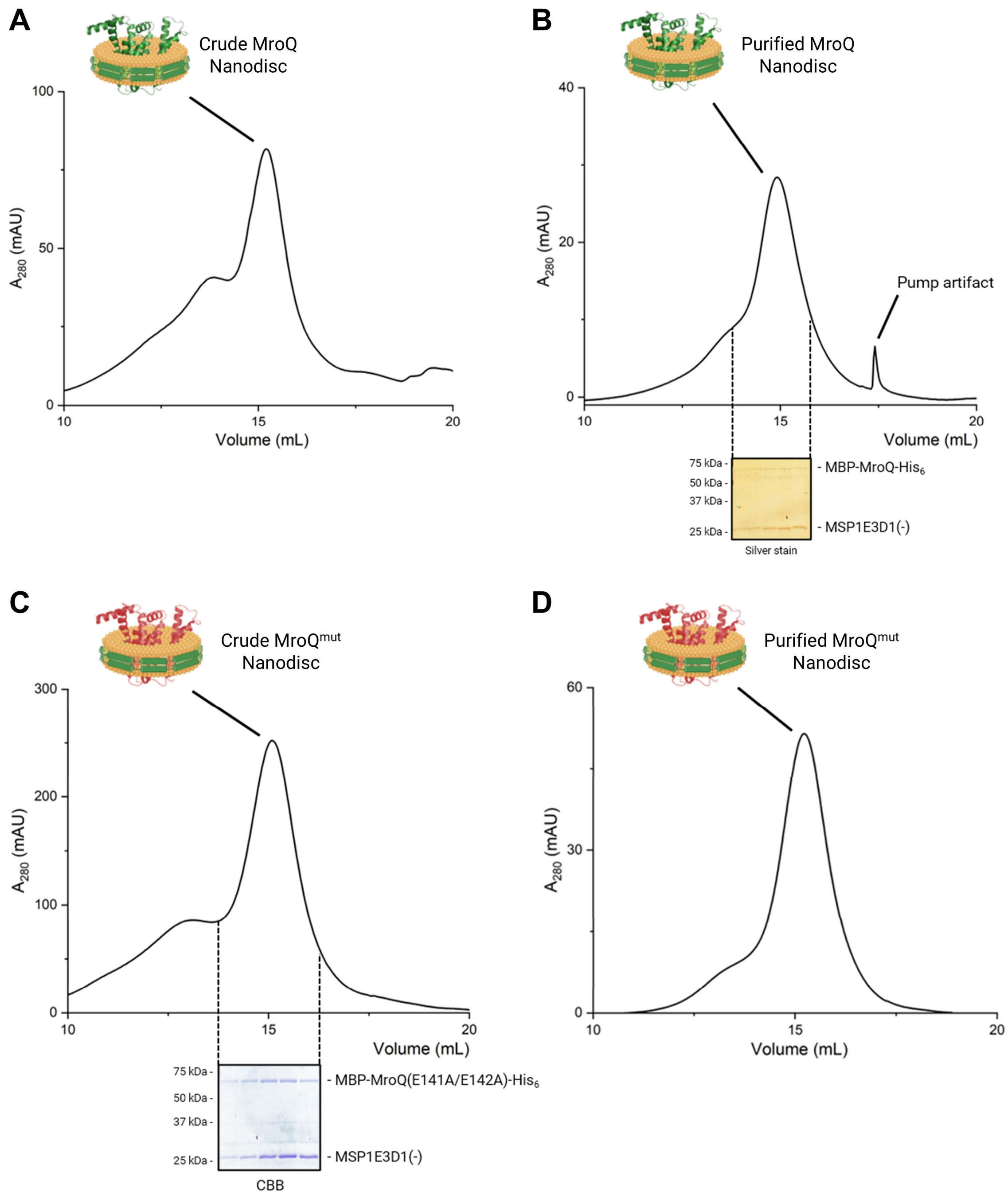


**Figure S6.** SEC and SDS-PAGE characterization of MroQ nanodiscs. (*A* and *B*) Characterization of crude (*A*) and purified (*B*) nanodiscs harboring MBP-MroQ-$His_6$. (*C* and *D*) Characterization of crude (*C*) and purified (*D*) nanodiscs harboring MBP-$MroQ^{mut}$-$His_6$. $MroQ^{mut}$ refers to the inactive E141A/E142A double mutant. A Superose 6 Increase 10/300 GL was used for SEC purification and analysis. MSP1E3D1(-) refers to MSP1E3D1 that has been cleaved with TEV protease for $His_7$-tag removal. In (*C*), CBB stands for Coomassie Brilliant Blue. All MroQ structures shown are AlphaFold 3 models (10).

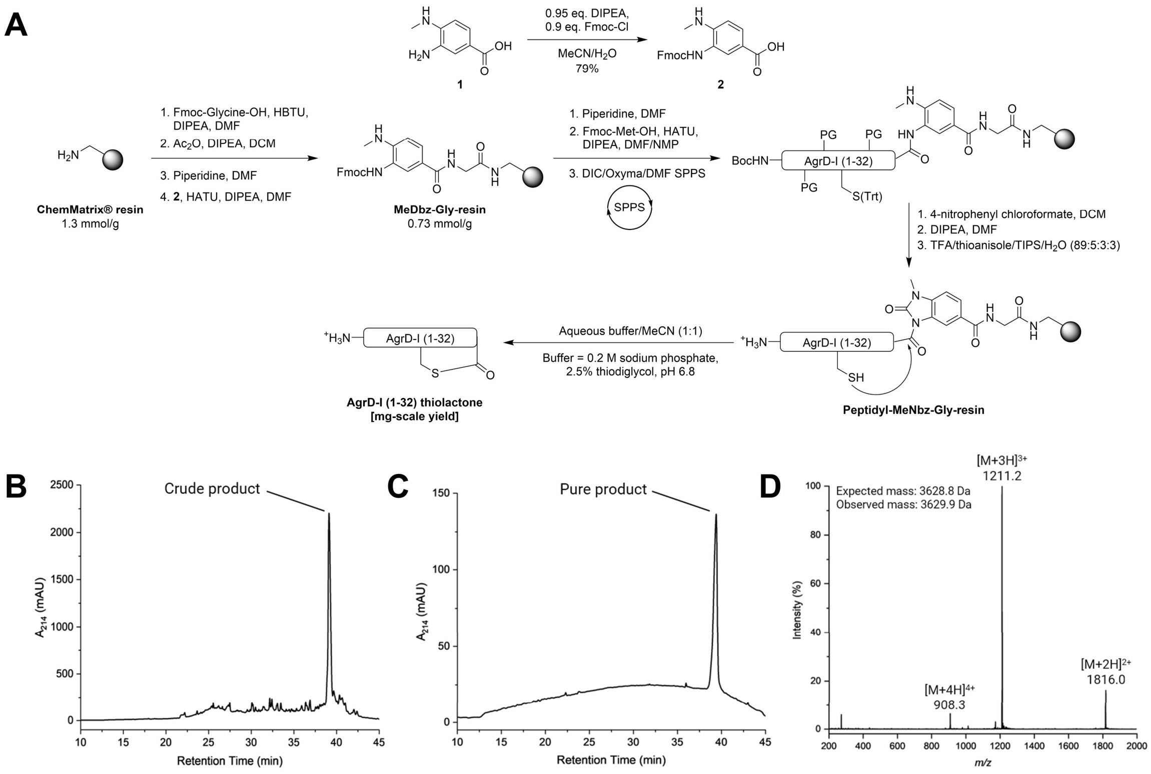


**Figure S7.** Synthesis of AgrD-I (1-32) thiolactone. (*A*) Reaction scheme showing how AgrD-I (1-32) thiolactone was synthetically produced using a second-generation *N*-acylurea Dawson linker. (*B* and *C*) Analytical $C_{18}$ RP-HPLC traces of crude (*B*) and purified (*C*) AgrD-I (1-32) thiolactone. The gradient was 5-100% solvent B over 35 min at 50°C, where solvent A was water + 0.1% TFA and solvent B was 10% water + 90% acetonitrile + 0.1% TFA. (*D*) Mass spectrum of the purified product.

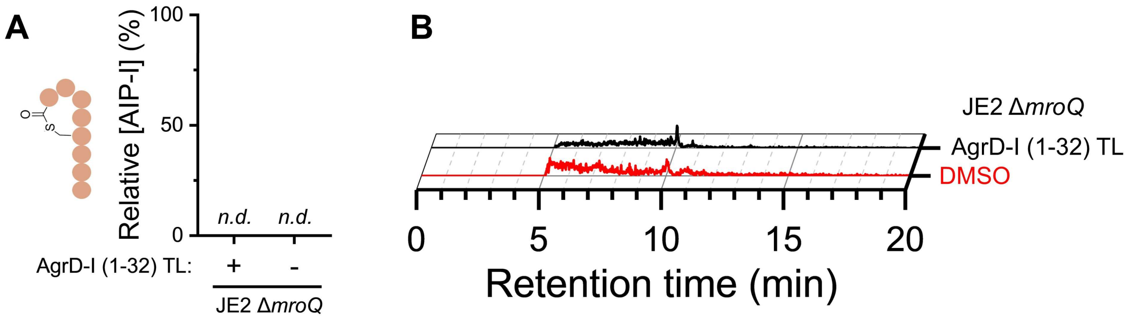


**Figure S8.** Cleavage of synthetic AgrD-I (1-32) thiolactone is dependent on MroQ. (*A*) AIP-I quantification of JE2 Δ*mroQ* cells grown in media containing a spike-in of either synthetic AgrD-I (1-32) thiolactone (+) or DMSO (-). Data is presented as mean ± SD (*n* = 3 biological replicates), normalized by an internal standard, and reported relative to WT cells treated with synthetic AgrD-I (1-32) thiolactone (*n.d.* means not detected). (*B*) Representative normalized EIC traces used for quantification in (*A*).

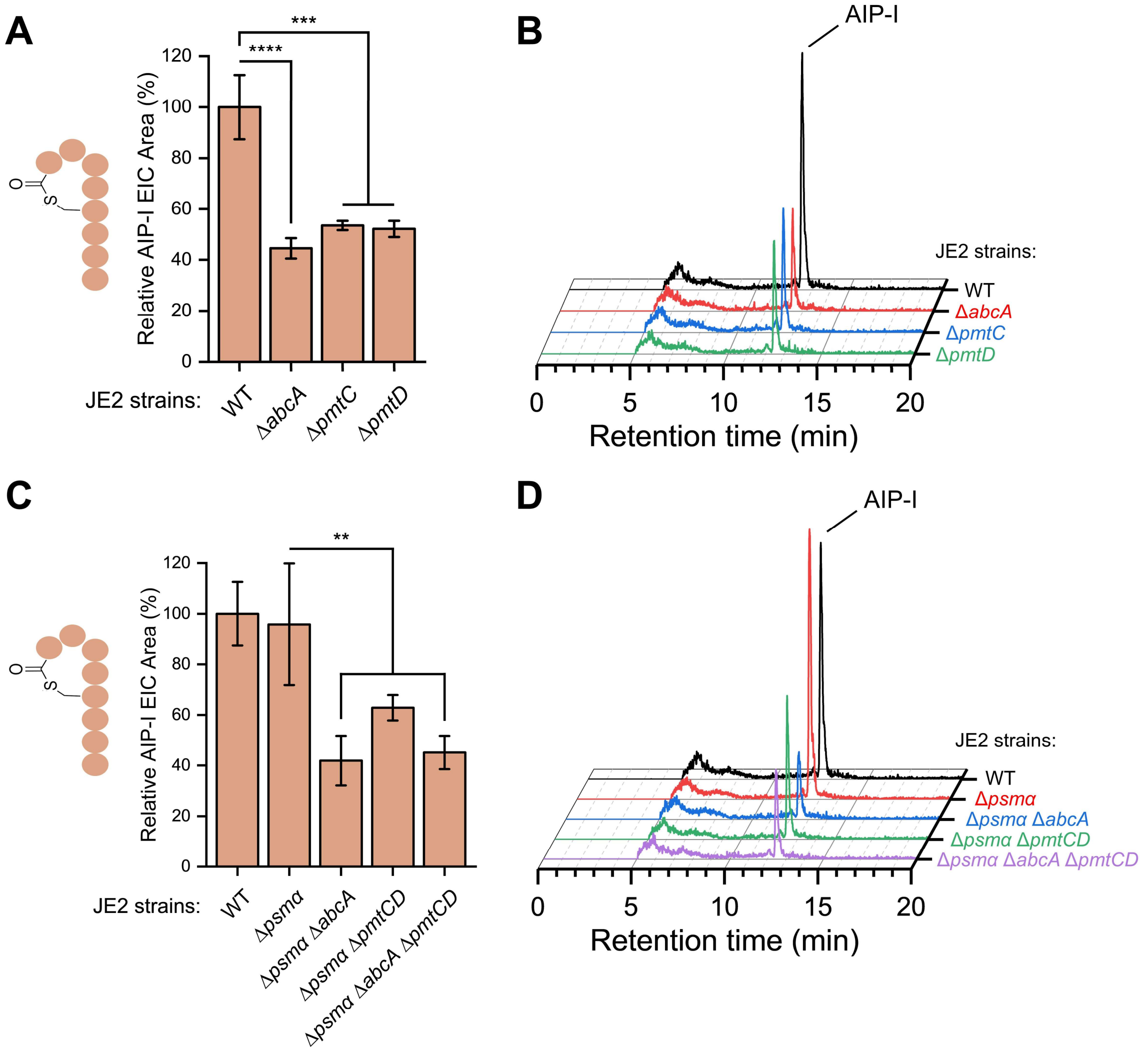


**Figure S9.** Gene deletion of *abcA* and *pmtCD* in JE2 reduces AIP-I production. (*A*) Quantification of AIP-I in the conditioned media derived from the indicated JE2 cultures grown for 18 hrs. Interestingly, we found that Δ*pmtC* and Δ*pmtD* strains of JE2 are viable, even though the *pmt* operon is essential (11). (*B*) Representative normalized EIC traces used for quantification in (*A*). (*C*) Quantification of AIP-I in the conditioned media derived from the indicated JE2 cultures grown for 18 hrs. (*D*) Representative normalized EIC traces used for quantification in (*C*). For (*A*) and (*C*), data is presented as mean ± SD ($n$ = 3 biological replicates), normalized by $OD_{600}$, and reported relative to WT samples. Statistical analysis is by 1-way ANOVA and Tukey HSD post-hoc analysis (****, $P < 0.0001$; ***, $P < 0.001$; **, $P < 0.01$).

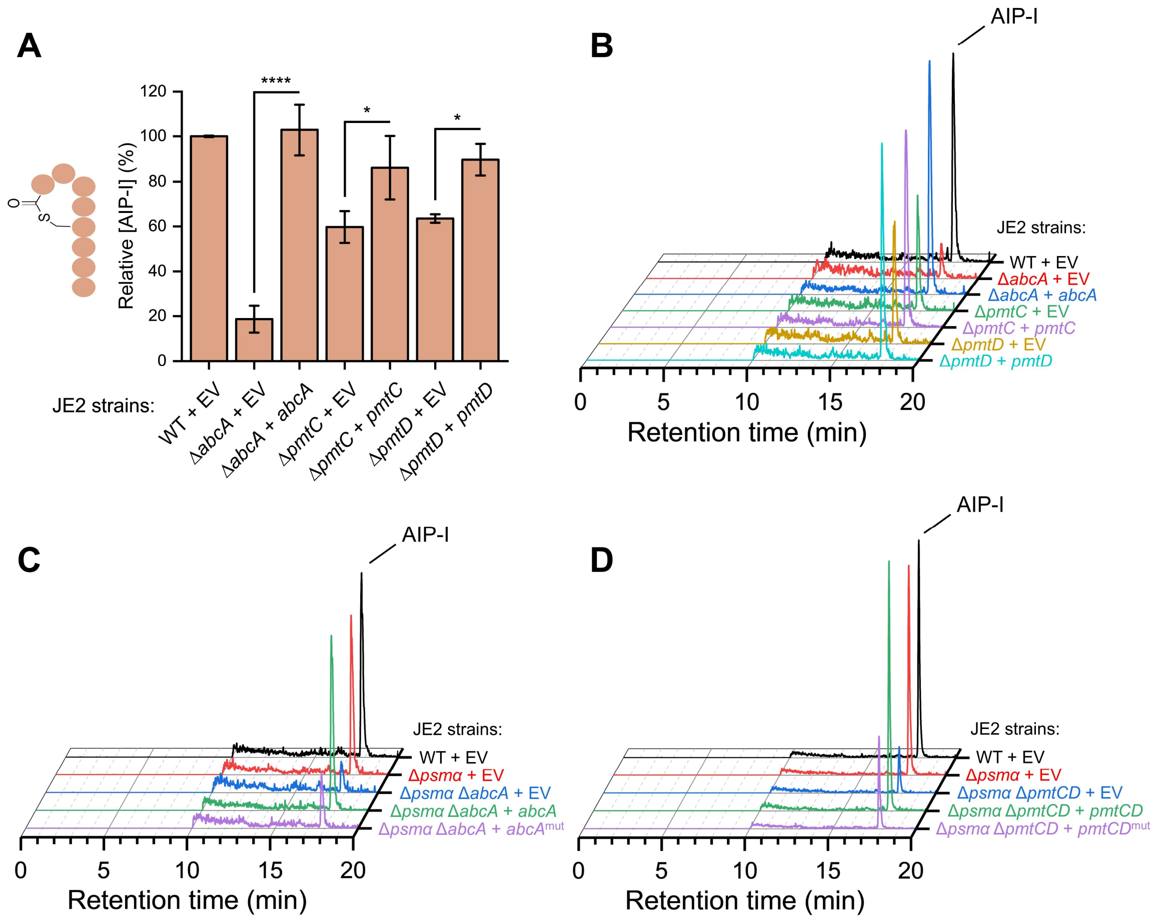


**Figure S10.** Genetic complementation with ATPase-active AbcA and PmtCD rescues AIP-I production in JE2 *abcA*/*pmtCD* knockout strains. (*A*) Quantification of AIP-I in the conditioned media derived from the indicated JE2 strains harboring complementation plasmids that utilize the native *abcA* promoter. Cultures were grown for 12 hrs. Data is presented as mean ± SD ($n$ = 3 biological replicates), normalized using both an internal standard and $OD_{600}$, and reported relative to WT samples. Statistical analysis is by 1-way ANOVA and Tukey HSD post-hoc analysis (****, $P < 0.0001$; *, $P < 0.05$). (*B*) Representative normalized EIC traces used for quantification in (*A*). (*C*) Representative normalized EIC traces used for quantification in Fig. 5*A*. (*D*) Representative normalized EIC traces used for quantification in Fig. 5*B*. EV refers to the empty vector control. Transporters with inactive ATPase domains are referred to as either *abcA*$^{mut}$ (K374M mutant) or *pmtCD*$^{mut}$ (E145Q mutant in PmtC).

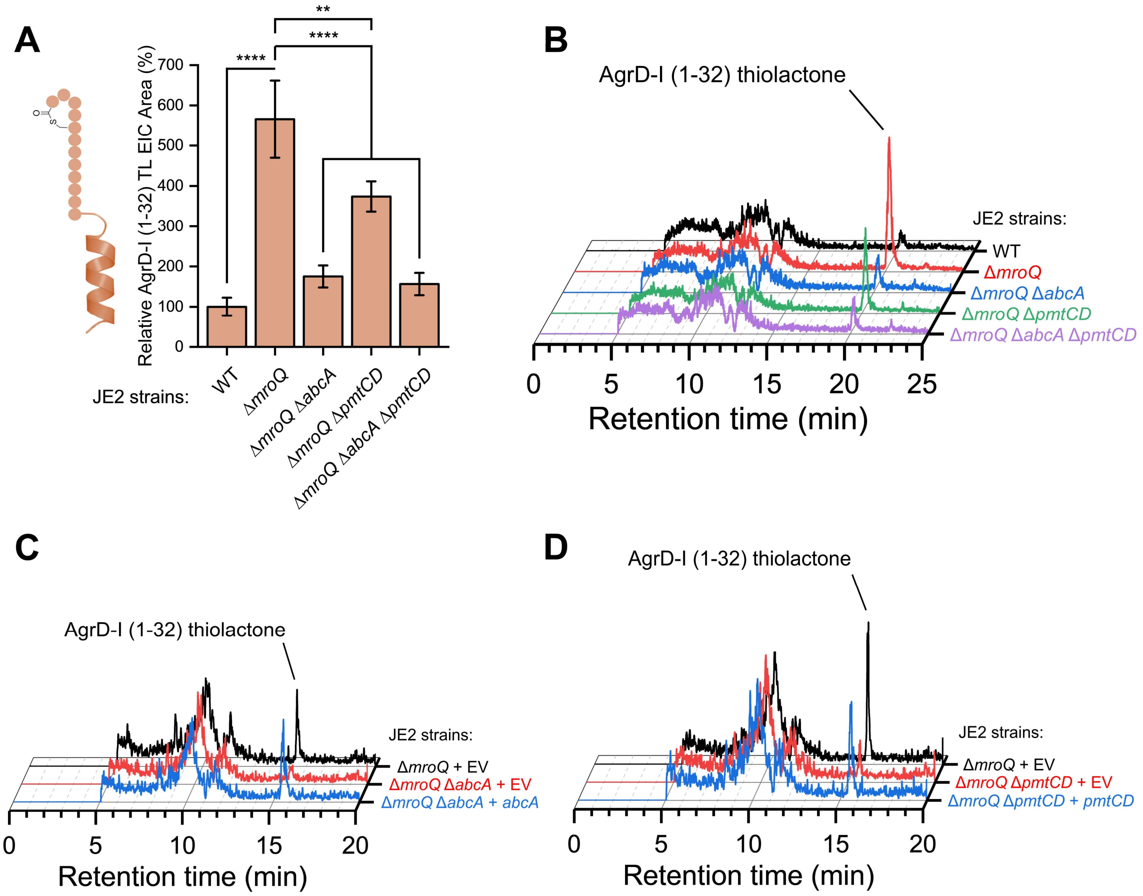


**Figure S11.** AbcA and PmtCD are involved in the export of AgrD-I (1-32) thiolactone. (*A*) Quantification of AgrD-I (1-32) thiolactone in the conditioned media derived from the indicated JE2 cultures grown for 18 hrs. Data is presented as mean ± SD ($n$ = 3 biological replicates), normalized using $OD_{600}$, and reported relative to WT samples. Statistical analysis is by 1-way ANOVA and Tukey HSD post-hoc analysis (****, $P < 0.0001$; **, $P < 0.01$). (*B*) Representative normalized EIC traces used for quantification in (*A*). (*C*) Representative normalized EIC traces used for quantification in Fig. 5*C*. (*D*) Representative normalized EIC traces used for quantification in Fig. 5*D*. EV refers to the empty vector control.

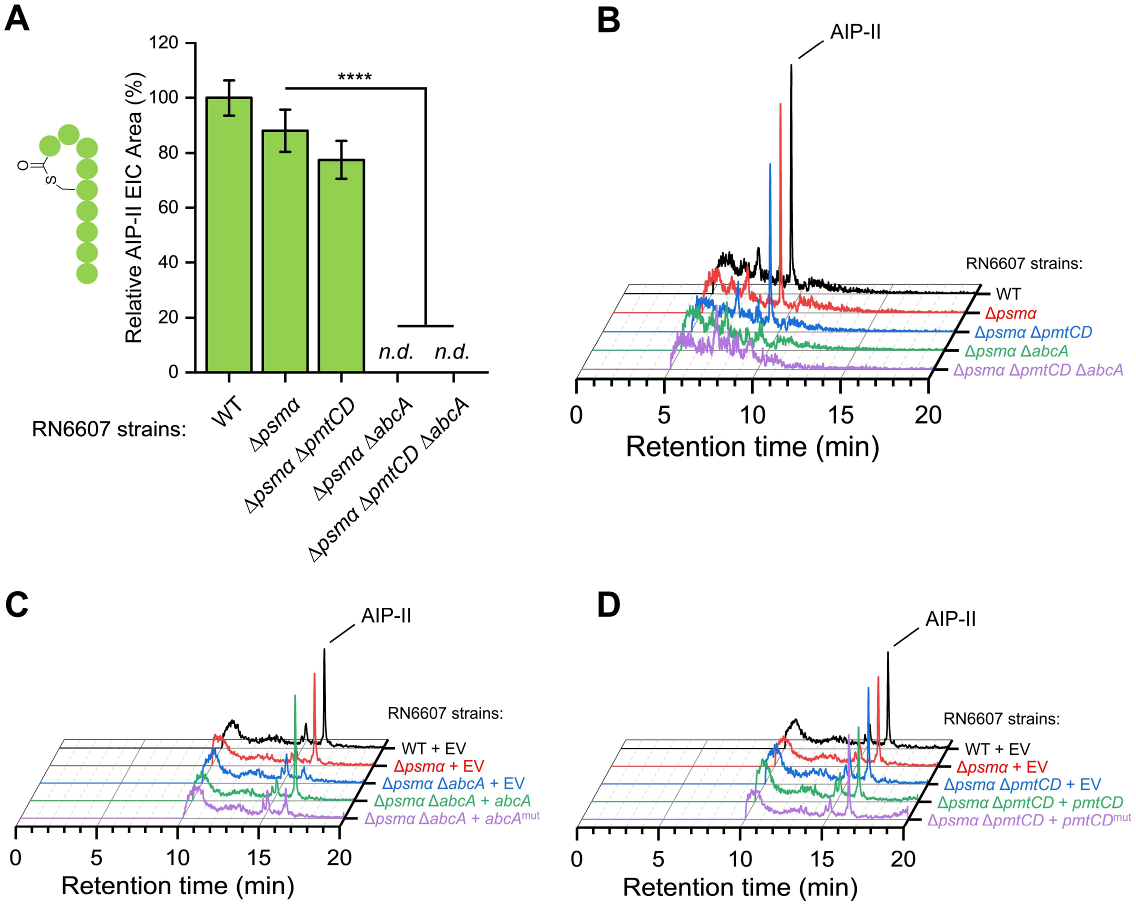


**Figure S12.** Genetic complementation with ATPase-active AbcA rescues AIP-II production in RN6607 *abcA* knockout strains. (*A*) Quantification of AIP-II in the conditioned media derived from the indicated RN6607 cultures grown for 18 hrs. Data is presented as mean ± SD ($n$ = 3 biological replicates), normalized using $OD_{600}$, and reported relative to WT samples. Statistical analysis is by 1-way ANOVA and Tukey HSD post-hoc analysis (****, $P < 0.0001$; *n.d.* means not detected). (*B*) Representative normalized EIC traces used for quantification in (*A*). (*C*) Representative normalized EIC traces used for quantification in Fig. 5*E*. (*D*) Representative normalized EIC traces used for quantification in Fig. 5*F*. EV refers to the empty vector control. Transporters with inactive ATPase domains are referred to as either *abcA*$^{mut}$ (K374M mutant) or *pmtCD*$^{mut}$ (E145Q mutant in PmtC).

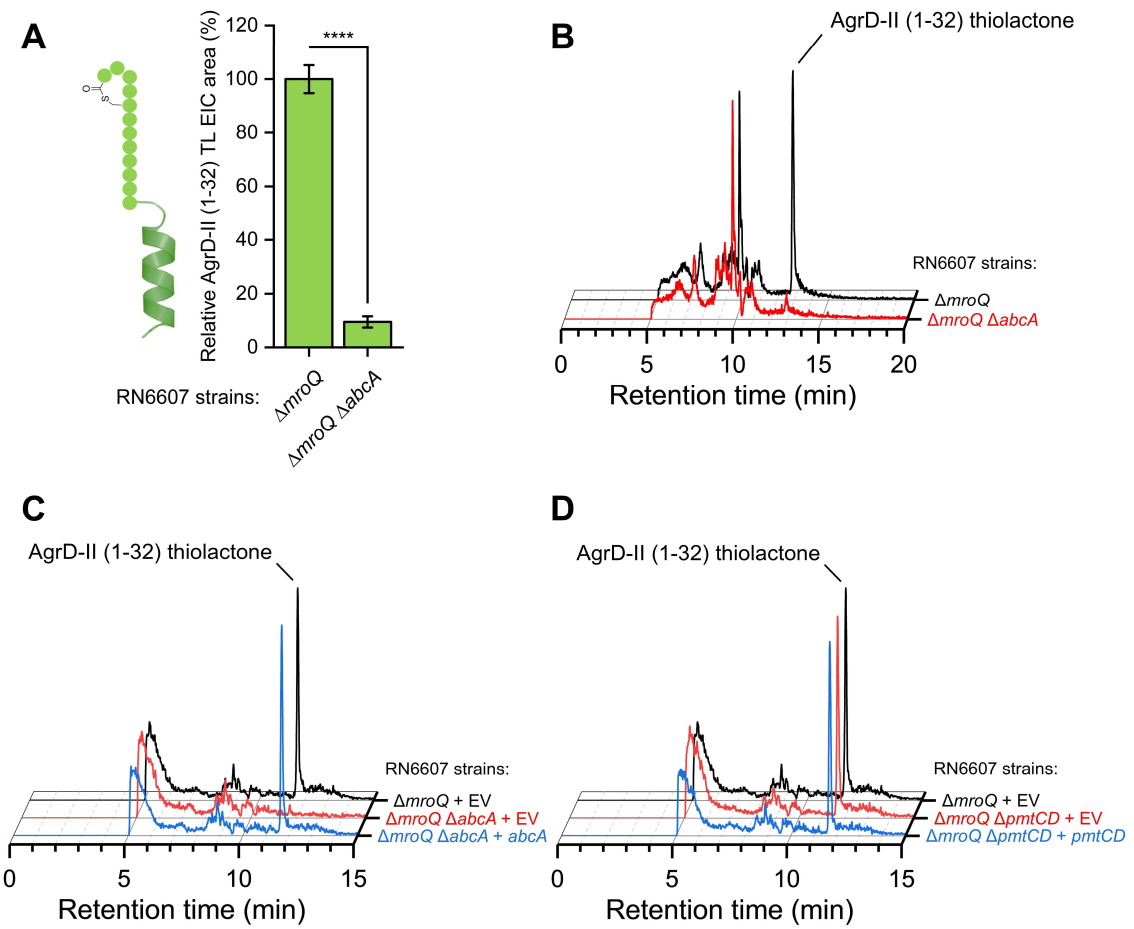


**Figure S13.** AbcA is involved in the export of AgrD-**II** (1-32) thiolactone. (*A*) Quantification of AgrD-**II** (1-32) thiolactone in the conditioned media derived from the indicated RN6607 cultures grown for 18 hrs. Data is presented as mean ± SD (*n* = 3 biological replicates), normalized using $OD_{600}$, and reported relative to Δ*mroQ* samples. Statistical analysis is by two-sample *t*-test (****, *P* < 0.0001). (*B*) Representative normalized EIC traces used for quantification in (*A*). (*C*) Representative normalized EIC traces used for quantification in Fig. 5*G*. (*D*) Representative normalized EIC traces used for quantification in Fig. 5*H*. EV refers to the empty vector control.

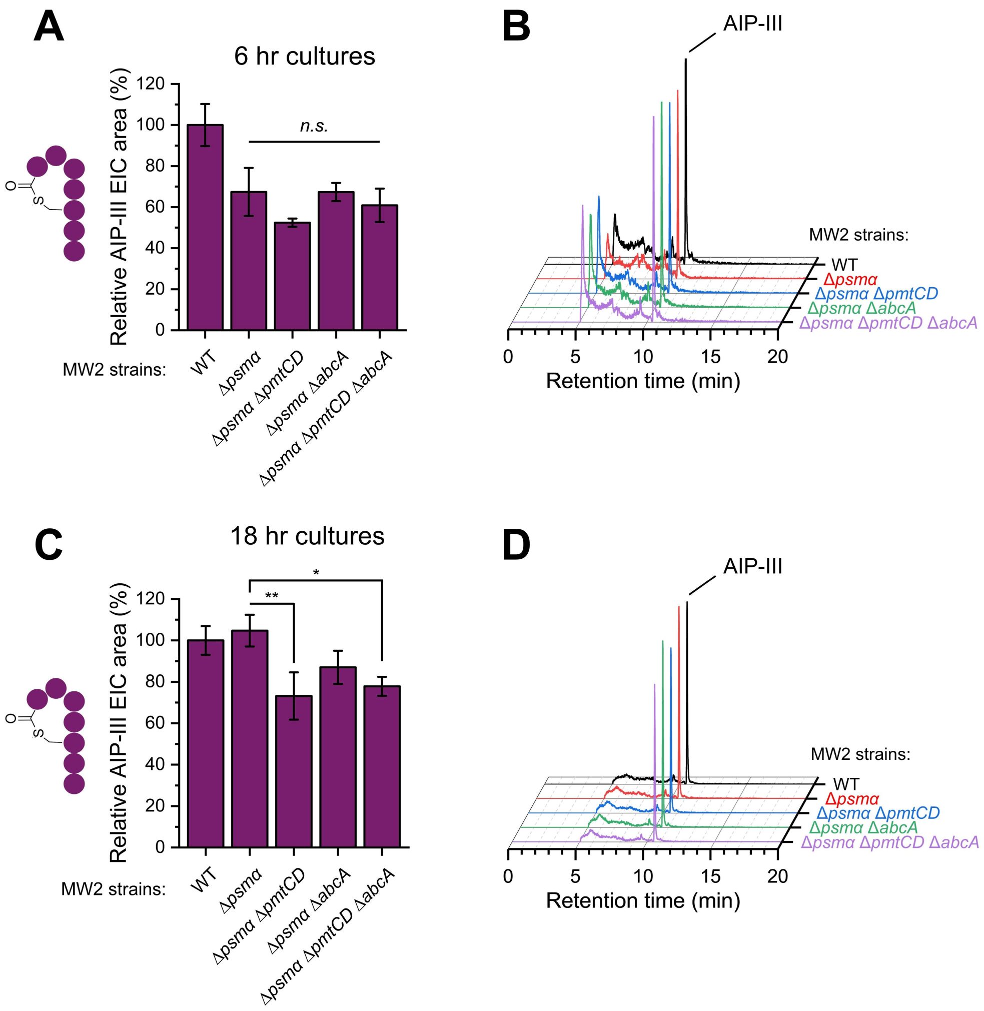


**Figure S14.** AbcA and PmtCD are not important for AIP-III biosynthesis. (*A* and *C*) Quantification of AIP-III in the conditioned media derived from the indicated MW2 cultures grown for 6 hrs (*A*) and 18 hrs (*C*). Data is presented as mean ± SD ($n$ = 3 biological replicates), normalized using $OD_{600}$, and reported relative to WT samples. Statistical analysis is by 1-way ANOVA and Tukey HSD post-hoc analysis (**, $P < 0.01$; *, $P < 0.05$; *n.s.* means not significant). (*B*) Representative normalized EIC traces used for quantification in (*A*). (*D*) Representative normalized EIC traces used for quantification in (*C*).

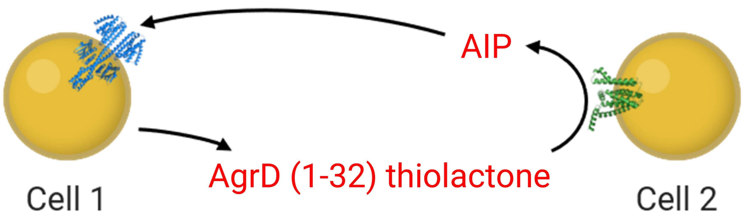


**Figure S15.** A model depicting AgrD (1-32) thiolactone as a kind of “chemical radar” that *S. aureus* can use for paracrine detection. AgrC (blue) and MroQ (green) structures are AlphaFold 3 models (10).

**Tables**

**Table S1.** Strains used in this study.

| Strain(s) | Description | Origin |
|---|---|---|
| ***E. coli* strains** | | |
| DH5α | Cloning strain for plasmid manipulation | Invitrogen |
| Stbl3 | Cloning strain for plasmid manipulation | Invitrogen |
| BL21(DE3) | Protein expression strain | Invitrogen |
| C43(DE3) | Membrane protein expression strain | Invitrogen |
| BL21(DE3)-RIL | Strain used for TEV protease expression | Addgene |
| ***S. aureus* strains** | | |
| RN4220 | Restriction-deficient *S. aureus* host strain | Kreiswirth *et al*. 1983 (12) |
| RN9222 | Group-I *agr* reporter strain for AIP-I detection | Lyon *et al*. 2000 (13) |
| NTML array | Library of JE2 Tn mutants | BEI resources |
| JE2 WT | *agr* group-I background strain for the NTML array | Fey *et al.* 2013 (14) |
| JE2 Δ*abcA* | *abcA* gene deletion mutant of JE2 | This study |
| JE2 Δ*pmtC* | *pmtC* gene deletion mutant of JE2 | This study |
| JE2 Δ*pmtD* | *pmtD* gene deletion mutant of JE2 | This study |
| JE2 Δ*psmα* | *psmα* gene deletion mutant of JE2 | This study |
| JE2 Δ*psmα* Δ*abcA* | *psmα*/*abcA* double gene deletion mutant of JE2 | This study |
| JE2 Δ*psmα* Δ*pmtCD* | *psmα*/*pmtCD* double gene deletion mutant of JE2 | This study |
| JE2 Δ*psmα* Δ*abcA* Δ*pmtCD* | *psmα*/*abcA*/*pmtCD* triple gene deletion mutant of JE2 | This study |
| JE2 Δ*mroQ* | *mroQ* gene deletion mutant of JE2 | This study |

| | | |
|---|---|---|
| JE2 Δ*mroQ* Δ*abcA* | *mroQ*/*abcA* double gene deletion mutant of JE2 | This study |
| JE2 Δ*mroQ* Δ*pmtCD* | *mroQ*/*pmtCD* double gene deletion mutant of JE2 | This study |
| JE2 Δ*mroQ* Δ*abcA* Δ*pmtCD* | *mroQ*/*abcA*/*pmtCD* triple gene deletion mutant of JE2 | This study |
| RN6607 WT | *agr* group-II strain | Ji *et al*. 1997 (15) |
| RN6607 Δ*psmα* | *psmα* gene deletion mutant of RN6607 | This study |
| RN6607 Δ*psmα* Δ*abcA* | *psmα*/*abcA* double gene deletion mutant of RN6607 | This study |
| RN6607 Δ*psmα* Δ*pmtCD* | *psmα*/*pmtCD* double gene deletion mutant of RN6607 | This study |
| RN6607 Δ*psmα* Δ*pmtCD* Δ*abcA* | *psmα*/*abcA*/*pmtCD* triple gene deletion mutant of RN6607 | This study |
| RN6607 Δ*mroQ* | *mroQ* gene deletion mutant of RN6607 | Zhao *et al*. 2022 (7) |
| RN6607 Δ*mroQ* Δ*abcA* | *mroQ*/*abcA* double gene deletion mutant of RN6607 | This study |
| RN6607 Δ*mroQ* Δ*pmtCD* | *mroQ*/*pmtCD* double gene deletion mutant of RN6607 | This study |
| MW2 WT | *agr* group-III strain | CDC, 1999 (16) |
| MW2 Δ*psmα* | *psmα* gene deletion mutant of MW2 | This study |
| MW2 Δ*psmα* Δ*abcA* | *psmα*/*abcA* double gene deletion mutant of MW2 | This study |
| MW2 Δ*psmα* Δ*pmtCD* | *psmα*/*pmtCD* double gene deletion mutant of MW2 | This study |
| MW2 Δ*psmα* Δ*pmtCD* Δ*abcA* | *psmα*/*abcA*/*pmtCD* triple gene deletion mutant of MW2 | This study |
| RN4850 WT | *agr* group-IV strain | BEI resources |
| RN4850 Δ*psmα* | *psmα* gene deletion mutant of RN4850 | This study |
| RN4850 Δ*mroQ* | *mroQ* gene deletion mutant of RN4850 | This study |

**Table S2.** Plasmids used in this study.

| Plasmid name | Description | Origin |
|---|---|---|
| **Protein Expression** | | |
| pMSP1E3D1 | Bacterial expression of an extended membrane scaffold protein (MSP1E3D1) for nanodisc formation | Denisov *et al*. 2007 (17) |
| pMroQ | Bacterial expression of MBP-MroQ-$His_6$ | Zhao *et al*. 2022 (7) |
| pMroQ$^{mut}$ | Bacterial expression of MBP-MroQ(E141A/E142A)-$His_6$ | Zhao *et al*. 2022 (7) |
| pRK793 | Bacterial expression of $His_6$-tagged TEV protease | Kapust *et al*. 2001 (18) |
| **Gene Deletion** | | |
| pMAD | Parent shuttle vector for *S. aureus* allelic exchange | Arnaud *et al*. 2004 (1) |
| pMAD_JE2_psmα | Plasmid used to delete *psmα* from the JE2 genome | This study |
| pMAD_JE2_mroQ | Plasmid used to delete *mroQ* from the JE2 genome | This study |
| pMAD_JE2_abcA | Plasmid used to delete *abcA* from the JE2 genome | This study |
| pMAD_JE2_pmtC | Plasmid used to delete *pmtC* from the JE2 genome | This study |
| pMAD_JE2_pmtD | Plasmid used to delete *pmtD* from the JE2 genome | This study |
| pMAD_JE2_pmtCD | Plasmid used to delete *pmtCD* from the JE2 genome | This study |
| pMAD_RN6607_psmα | Plasmid used to delete *psmα* from the RN6607 genome | This study |
| pMAD_RN6607_abcA | Plasmid used to delete *abcA* from the RN6607 genome | This study |
| pMAD_RN6607_pmtCD | Plasmid used to delete *pmtCD* from the RN6607 genome | This study |
| pMAD_MW2_psmα | Plasmid used to delete *psmα* from the MW2 genome | This study |
| pMAD_MW2_abcA | Plasmid used to delete *abcA* from the MW2 genome | This study |
| pMAD_MW2_pmtCD | Plasmid used to delete *pmtCD* from the MW2 genome | This study |
| pMAD_RN4850_psmα | Plasmid used to delete *psmα* from the RN4850 genome | This study |

| | | |
|---|---|---|
| pMAD_RN4850_mroQ | Plasmid used to delete *mroQ* from the RN4850 genome | This study |
| **Genetic complementation** | | |
| pCN51_EV | Empty vector containing a cadmium-inducible promoter | Charpentier *et al*. 2004 (2) |
| pCN51_pmtCD | Cadmium-inducible PmtCD expression | This study |
| pCN51_pmtCD$^{mut}$ | Cadmium-inducible PmtC(E145Q)D expression | This study |
| pOS1_sGFP-P$_{sarA}$-*sod* RBS | *sarA* promoter controlling expression of *sgfp* in a pOS1(Cm$^{r}$) vector | Benson *et al*. 2012 (3) |
| pOS1(Em$^{r}$)_EV | A modified version of pOS1_sGFP-P$_{sarA}$-*sod* RBS in which Cm$^{r}$ is replaced with Em$^{r}$, the sGFP ORF and protein A gene fragments are deleted, and the P$_{sarA}$ promoter is replaced by P$_{abcA}$ | This study |
| pOS1(Em$^{r}$)_abcA | P$_{abcA}$-driven AbcA expression | This study |
| pOS1(Em$^{r}$)_abcA$^{mut}$ | P$_{abcA}$-driven AbcA(K374M) expression | This study |
| pOS1(Em$^{r}$)_pmtC | P$_{abcA}$-driven PmtC expression | This study |
| pOS1(Em$^{r}$)_pmtD | P$_{abcA}$-driven PmtD expression | This study |
| pOS1(Em$^{r}$)_pmtCD | P$_{abcA}$-driven PmtCD expression | This study |
| pOS1(Em$^{r}$)_pmtCD$^{mut}$ | P$_{abcA}$-driven PmtC(E145Q)D expression | This study |

**Table S3.** Primers used in this study.

| Primer name | Description | Sequence |
|---|---|---|
| **pMAD construction** | | |
| JE2_psma_u_f | Fwd primer upstream of *psmα* in the JE2 (and MW2) genome | GATGCATGCCATGGGTTCGATCAAAAGGTATCGTATGGC |
| JE2_psma_u_r | Rev primer upstream of *psmα* in the JE2 (and RN6607/MW2/RN4850) genome | GCAAAGGAGGTAATCTTATTTAAGCGAATTGAATACTTAAAATTCTCAGG |
| JE2_psma_d_f | Fwd primer downstream of *psmα* in the JE2 (and RN6607/MW2/RN4850) genome | GTATTCAATTCGCTTAAATAAGATTACCTCCTTTGCTTATGAGTTAAC |
| JE2_psma_d_r | Rev primer downstream of *psmα* in the JE2 (and RN6607) genome | CGAGCTCCCGGGCATAAAAATGTCATGCTTGATAATTTCGCAG |
| JE2_mroQ_u_f | Fwd primer upstream of *mroQ* in the JE2 genome | ATGGTACCCGGGCAATTTCTTCATCTGCTGCTGAAATC |
| JE2_mroQ_u_r | Rev primer upstream of *mroQ* in the JE2 genome | GTGATTCGTTTTTTTTATTAAATTTTCCTCCAAATATTTGTTTATAATTTATTTTATCG |
| JE2_mroQ_d_f | Fwd primer downstream of *mroQ* in the JE2 genome | CAAATATTTGGAGGAAAATTTAATAAAAAAAACGAATCACACACGG |
| JE2_mroQ_d_r | Rev primer downstream of *mroQ* in the JE2 genome | ACGTCGACGCGTCTCAAACAACCAAAACCCAAGTAC |
| JE2_abcA_u_f | Fwd primer upstream of *abcA* in the JE2 genome | ATGGTACCCGGGCTTTGTTGGTGCAAATGTACGTAATC |
| JE2_abcA_u_r | Rev primer upstream of *abcA* in the JE2 genome | CTTATATATATAAAATTAAGTTTTACCTCTTCTGTTTGAAATTTATAGTTAATTC |
| JE2_abcA_d_f | Fwd primer downstream of *abcA* in the JE2 genome | CAAACAGAAGAGGTAAAACTTAATTTTATATATATAAGTAAGCTTGGAGCAAATACACATATAC |
| JE2_abcA_d_r | Rev primer downstream of *abcA* in the JE2 (and RN6607/MW2) genome | ACGTCGACGCGTCTGGATTCAACAAGCATGACAC |
| JE2_pmtC_u_f | Fwd primer upstream of *pmtC* in the JE2 genome | ATGGTACCCGGGGGGGACCAAGAGCAAATGAAG |
| JE2_pmtC_u_r | Rev primer upstream of *pmtC* in the JE2 genome | GGAGGCAGTCGATGAATGAGAATTTTAAATTTAGTTAAGTATGATTTTTATAGTATATTTAAAAG |
| JE2_pmtC_d_f | Fwd primer downstream of *pmtC* in the JE2 genome | CTAAATTTAAAATTCTCATTCATCGACTGCCTCCTTTAAATTTTC |
| JE2_pmtC_d_r | Rev primer downstream of *pmtC* in the JE2 genome | GATATCGGATCCGAAAAAATAGCTGATTATATCATTCACTTAAGTGAC |

| | | |
|---|---|---|
| JE2_pmtD_u_f | Fwd primer upstream of *pmtD* in the JE2 genome | TGCATGCCATGGCTGTTATAAAATTTAAAACGATCAAACTTAATTTTTTTATTTAC |
| JE2_pmtD_u_r | Rev primer upstream of *pmtD* in the JE2 genome | CAAAGAGGTGATAAATAATAATAAAAATAATTTTGAGGTTGGGAATTTTAAATTTTC |
| JE2_pmtD_d_f | Fwd primer downstream of *pmtD* in the JE2 genome | CCTCAAAATTATTTTTATTATTATTTATCACCTCTTTGATTTATATTGAAGTACG |
| JE2_pmtD_d_r | Rev primer downstream of *pmtD* in the JE2 genome | CGAGCTCCCGGGCATTCCCAGATTTATATTCATATATTTTAAATATTGGTTTTC |
| JE2_pmtCD_u_f | Fwd primer upstream of *pmtCD* in the JE2 (and RN6607/MW2) genome | TGCATGCCATGGCTGTTATAAAATTTAAAACGATCAAACTTAATTTTTTTATTTAC |
| JE2_pmtCD_u_r | Rev primer upstream of *pmtCD* in the JE2 (and RN6607/MW2) genome | TTAAAGGAGGCAGTCGATGATAATAAAAATAATTTTGAGGTTGGGAATTTTAAATTTTC |
| JE2_pmtCD_d_f | Fwd primer downstream of *pmtCD* in the JE2 (and RN6607/MW2) genome | CCTCAAAATTATTTTTATTATCATCGACTGCCTCCTTTAAATTTTC |
| JE2_pmtCD_d_r | Rev primer downstream of *pmtCD* in the JE2 (and RN6607/MW2) genome | CGAGCTCCCGGGGAAAAAATAGCTGATTATATCATTCACTTAAGTGAC |
| RN6607_psma_u_f | Fwd primer upstream of *psmα* in the RN6607 genome | GATGCATGCCATGGGTTCGATCAAAAGGTATCGTATGGTTAG |
| RN6607_psma_f_gib | Fwd Gibson primer upstream of insert in pMAD_RN6607_psmα | CATTAACTAGACAGATCTATCGATGCATGCCATGGGTTCG |
| RN6607_psma_r_gib | Rev Gibson primer downstream of insert in pMAD_RN6607_psmα | GCAGAAGCTTCTAGAATTCGAGCTCCCGGGCA |
| RN6607_abcA_u_f | Fwd primer upstream of *psmα* in the RN6607 (and MW2) genome | CATGGTACCCGGGCTTTGTTGGTGCAAATGAACGTAATC |
| RN6607_abcA_u_r | Rev primer upstream of *abcA* in the RN6607 (and MW2) genome | GCTCCAAGCTTACTTATATATAAAAAGTTTTACCTCTTCTGTTTGAAATTTATAGTTAATTC |
| RN6607_abcA_d_f | Fwd primer downstream of *abcA* in the RN6607 (and MW2) genome | CAAACAGAAGAGGTAAAACTTTTTATATATAAGTAAGCTTGGAGCAAATACAC |
| RN6607_abcA_f_gib | Fwd Gibson primer upstream of insert in pMAD_RN6607_abcA (and pMAD_MW2_abcA) | GATCTATCGATGCATGCCATGGTACCCGGGCTTTG |
| RN6607_abcA_r_gib | Rev Gibson primer downstream of insert in pMAD_RN6607_abcA (and pMAD_MW2_abcA) | GATATCGGATCCATATGACGTCGACGCGTCTGG |
| RN6607_pmtCD_f_gib | Fwd Gibson primer upstream of insert in pMAD_RN6607_pmtCD (and pMAD_MW2_pmtCD) | CATTAACTAGACAGATCTATCGATGCATGCCATGGCTGTTATAA |

| | | |
|---|---|---|
| RN6607_pmtCD_r_gib | Rev Gibson primer downstream of insert in pMAD_RN6607_pmtCD (and pMAD_MW2_pmtCD) | GCAGAAGCTTCTAGAATTCGAGCTCCCGGGGAAA |
| MW2_psma_d_r | Rev primer downstream of *psmα* in the MW2 genome | CGAGCTCCCGGGCACATAAAAATGTCATACTTGATAATTTCGC |
| MW2_psma_f_gib | Fwd Gibson primer upstream of insert in pMAD_MW2_psmα | CATTAACTAGACAGATCTATCGATGCATGCCATGGGTTCG |
| MW2_psma_r_gib | Rev Gibson primer downstream of insert in pMAD_MW2_psmα | GCAGAAGCTTCTAGAATTCGAGCTCCCGGGCAC |
| RN4850_psma_u_f | Fwd primer upstream of *psmα* in the RN4850 genome | CATGGTACCCGGGGTTCGATCAAAAGGTATCGTATGGC |
| RN4850_psma_d_r | Rev primer downstream of *psmα* in the RN4850 genome | GACGTCGACGCGTCATAAAAATGTCATGCTTGATAATTTCGC |
| RN4850_psma_f_gib | Fwd Gibson primer upstream of insert in pMAD_RN4850_psmα | GATCTATCGATGCATGCCATGGTACCCGGGGTTCG |
| RN4850_psma_r_gib | Rev Gibson primer downstream of insert in pMAD_RN4850_psmα | GATATCGGATCCATATGACGTCGACGCGTCATAAA |
| RN4850_mroQ_u_f | Fwd primer upstream of *mroQ* in the RN4850 genome | CATGGTACCCGGGCAATTTCTTCATCTGCTGCAGAAATC |
| RN4850_mroQ_u_r | Rev primer upstream of *mroQ* in the RN4850 genome | GTGATTCGTTTTTTATAAATTTTCCTCCAAATATTTGTTTATAATTTATTTTATCG |
| RN4850_mroQ_d_f | Fwd primer downstream of *mroQ* in the RN4850 genome | CAAATATTTGGAGGAAAATTTATAAAAAACGAATCACACACGGATTATC |
| RN4850_mroQ_d_r | Rev primer downstream of *mroQ* in the RN4850 genome | GACGTCGACGCGTGAAATAAATAATAATGAGACTGGGACACCTC |
| RN4850_mroQ_f_gib | Fwd Gibson primer upstream of insert in pMAD_RN4850_mroQ | CTATCGATGCATGCCATGGTACCCGGGCAATTTC |
| RN4850_mroQ_r_gib | Rev Gibson primer downstream of insert in pMAD_RN4850_mroQ | GATATCGGATCCATATGACGTCGACGCGTGAAATAAATAATAATGAGACTG |
| **Complementation vector construction** | | |
| pCN51_pmtCD_f | Fwd primer to amplify *pmtCD* from the JE2 genome and *pmtCD*$^{mut}$ from pOS1(Em$^{r}$)_pmtCD$^{mut}$ | AGGATCCCCGGGATGAAATTAGAACATATTACAAAAAAATACGGC |
| pCN51_pmtCD_r | Rev primer to amplify *pmtCD* from the JE2 genome and *pmtCD*$^{mut}$ from pOS1(Em$^{r}$)_pmtCD$^{mut}$ | GAGCTCGGTACCTTAAATGTCTTTTTTAGCGACATATAAGATATTCAAAATG |
| pmtC_E145Q_f | Fwd site-directed mutagenesis primer for PmtC(E145Q) | CTTGGATCAGCCTACAAATGGTATG |
| pmtC_E145Q_r | Rev site-directed mutagenesis primer for PmtC(E145Q) | CATACCATTTGTAGGCTGATCCAAGATTAAAAATTTAGGTTTATTC |

| | | |
|---|---|---|
| PabcA_EV_f | Fwd primer for deleting *abcA* from pOS1(Em$^r$)_abcA to generate pOS1(Em$^r$)_EV | CAAACAGAAGAGGTAAAACTTTTTATATATATAAGTAAGCTTGGAGC |
| PabcA_EV_f | Rev primer for deleting *abcA* from pOS1(Em$^r$)_abcA to generate pOS1(Em$^r$)_EV | AGTTTTACCTCTTCTGTTTGAAATTTATAG |
| Erm_f | Fwd Gibson primer for amplifying Em$^r$ for insertion into pOS1(Em$^r$)_EV | CACTTTAGATAAAAATTTAGGAGGCATATCAAAATTGAATGAGACATGCTACACCTC |
| Erm_r | Rev Gibson primer for amplifying Em$^r$ for insertion into pOS1(Em$^r$)_EV | GTAAAAAGTACAGTCGGCATTATCTCATAAAAACTGGTTTAAGCCGACTGC |
| pOS1_cat_f | Fwd Gibson primer for removing Cm$^r$ for generation of pOS1(Em$^r$)_EV | TATGAGATAATGCCGACTGTACTTTTTAC |
| pOS1_cat_r | Rev Gibson primer for removing Cm$^r$ for generation of pOS1(Em$^r$)_EV | TTGATATGCCTCCTAAATTTTTATCTAAAGTG |
| PabcA_vector_fwd | Fwd Gibson backbone primer for generating pOS1(Em$^r$)_abcA | GATTTTAGAGTCCACGGTACAGTATGGTGCACTCTCAGTACAATCTG |
| PabcA_vector_rev | Rev Gibson backbone primer for generating pOS1(Em$^r$)_abcA | GCATTATTTTTCAAATTAACTGCAGGCCTGGGGTGCCTAATGAG |
| PabcA_abcA_fwd | Fwd Gibson insert primer for generating pOS1(Em$^r$)_abcA | GCCTGCAGTTAATTTGAAAAATAATGC |
| PabcA_abcA_rev | Rev Gibson insert primer for generating pOS1(Em$^r$)_abcA | CTGTACCGTGGACTCTAAAATC |
| abcA_K374M_f | Fwd site-directed mutagenesis primer for AbcA(K374M) | GGTCTGGTATGAGTACGATATTTAATCTGATAGAACGTATG |
| abcA_K374M_r | Rev site-directed mutagenesis primer for AbcA(K374M) | CAGATTAAATATCGTACTCATACCAGACCCAGAAGGGC |
| PabcA_pmtCvec_fwd | Fwd Gibson backbone primer for generating pOS1(Em$^r$)_pmtC | CAATATAAATCAAAGAGGTGATAAATAATTTTATATATATAAGTAAGCTTGGAGC |
| PabcA_pmtCvec_rev | Rev Gibson backbone primer for generating pOS1(Em$^r$)_pmtC and pOS1(Em$^r$)_pmtCD | GTAATATGTTCTAATTTCATAGTTTTACCTCTTCTGTTTG |
| PabcA_pmtCins_fwd | Fwd Gibson insert primer for generating pOS1(Em$^r$)_pmtC and pOS1(Em$^r$)_pmtCD | CAGAAGAGGTAAAACTATGAAATTAGAACATATTACAAAAAAATACG |
| PabcA_pmtCins_rev | Rev Gibson insert primer for generating pOS1(Em$^r$)_pmtC | GCTTACTTATATATATAAAATTATTTATCACCTCTTTGATTTATATTG |
| PabcA_pmtCDvec_fwd | Fwd Gibson backbone primer for generating pOS1(Em$^r$)_pmtD and pOS1(Em$^r$)_pmtCD | CTTATATGTCGCTAAAAAAGACATTTAATTTTATATATATAAGTAAGCTTGGAGC |
| PabcA_pmtDvec_rev | Rev Gibson backbone primer for generating pOS1(Em$^r$)_pmtD | CTAAATTTAAAATTCTCATAGTTTTACCTCTTCTGTTTG |

| | | |
|---|---|---|
| PabcA_pmtDins_fwd | Fwd Gibson insert primer for generating pOS1(Em$^r$)_pmtD | CAGAAGAGGTAAAACTATG AGAATTTTAAATTTAGTTAA GTATGATTTTTATAG |
| PabcA_pmtDins_rev | Rev Gibson insert primer for generating pOS1(Em$^r$)_pmtD and pOS1(Em$^r$)_pmtCD | CTTATATATATAAAATTAAA TGTCTTTTTTAGCGACATAT AAG |

### SI References


1. M. Arnaud, A. Chastanet, M. Débarbouillé, New vector for efficient allelic replacement in naturally nontransformable, low-GC-content, gram-positive bacteria. *Appl. Environ. Microbiol.* **70**, 6887–6891 (2004).
2. E. Charpentier *et al.*, Novel cassette-based shuttle vector system for gram-positive bacteria. *Appl. Environ. Microbiol.* **70**, 6076–6085 (2004).
3. M. A. Benson, S. Lilo, T. Nygaard, J. M. Voyich, V. J. Torres, Rot and SaeRS cooperate to activate expression of the staphylococcal superantigen-like exoproteins. *J. Bacteriol.* **194**, 4355–4354 (2012).
4. T. K. Ritchie *et al.*, Reconstitution of membrane proteins in phospholipid bilayer nanodiscs. *Methods Enzymol.* **464**, 211–231 (2009).
5. B. Wang, "Reconstitution and mechanistic studies on the staphylococcal *agr* quorum sensing circuit," PhD thesis, Rockefeller University, New York, NY (2016).
6. J. E. Tropea, S. Cherry, D. S. Waugh, Expression and purification of soluble $His_6$-tagged TEV protease. *Methods Mol. Biol.* **498**, 297–307 (2009).
7. A. Zhao *et al.*, Reconstitution of the *S. aureus agr* quorum sensing pathway reveals a direct role for the integral membrane protease MroQ in pheromone biosynthesis. *Proc. Natl. Acad. Sci. U.S.A.* **119** (2022).
8. J. B. Blanco-Canosa, B. Nardone, F. Albericio, P. E. Dawson, Chemical protein synthesis using a second-generation *N*-acylurea linker for the preparation of peptide-thioester precursors. *J. Am. Chem. Soc.* **137**, 7197–7209 (2015).
9. B. H. Gless, C. A. Olsen, On-resin peptide cyclization using the 3-amino-4-(methylamino)benzoic acid MeDbz linker. *Methods Mol. Biol.* **2371**, 101–115 (2022).
10. J. Abramson *et al.*, Accurate structure prediction of biomolecular interactions with AlphaFold 3. *Nature.* **630** 493–500 (2024).
11. S. S. Chatterjee *et al.*, Essential *Staphylococcus aureus* toxin export system. *Nat. Med.* **19**, 364–367 (2013).
12. B. N. Kreiswirth, The toxic shock syndrome exotoxin structural gene is not detectably transmitted by a prophage. *Nature*. **305**, 709–712 (1983).
13. G. J. Lyon, P. Mayville, T. W. Muir, R. P. Novick, Rational design of a global inhibitor of the virulence response in *Staphylococcus aureus*, based in part on localization of the site of inhibition to the receptor-histidine kinase, AgrC. *Proc. Natl. Acad. Sci. U.S.A.* **97**, 13330–13335 (2000).
14. P. D. Fey *et al.*, A genetic resource for rapid and comprehensive phenotype screening of nonessential *Staphylococcus aureus* genes. *mBio*. **4** (2013).
15. G. Ji, R. Beavis, R. P. Novick, Bacterial interference caused by autoinducing peptide variants. *Science.* **276**, 2027–2030 (1997).
16. Centers for Disease Control and Prevention (CDC), Four pediatric deaths from community-acquired methicillin-resistant Staphylococcus aureus — Minnesota and North Dakota, 1997-1999. *Morb*. *Mortal*. *Wkly*. *Rep*. **48**, 707–710 (1999).
17. I. G. Denisov, B. J. Baas, Y. V. Grinkova, S. G. Sligar, Cooperativity in cytochrome P450 3A4: linkages in substrate binding, spin state, uncoupling, and product formation. *J. Biol. Chem.* **282**, 7066–7076 (2007).
18. R. B. Kapust *et al.*, Tobacco etch virus protease: mechanism of autolysis and rational design of stable mutants with wild-type catalytic proficiency. *Protein Eng.* **14**, 993–1000 (2001).
